# Polycomb establishes TAD-scale H3K27me3 mega-domains to safeguard neuronal identity

**DOI:** 10.64898/2026.08.10.743812

**Authors:** Natsumi Yamada, Chika Ichihara, Kazunori Hojo, Koji Oishi, Hiroki Sugishita, Yukiko Gotoh

## Abstract

During development, pluripotent stem cells generate diverse cell types through gene regulatory networks orchestrated by combinations of transcription factors (TFs). Following terminal differentiation, however, cellular identities become remarkably stable and resistant to TF-mediated perturbation, yet the mechanisms underlying this stability remain poorly understood. Here, we identify Polycomb repressive complexes (PRCs) as key regulators of neuronal identity maintenance. Although PRCs are well known for repressing the promoters of developmental genes during early cell fate specification, we unexpectedly find that PRC-mediated H3K27me3 expands into megabase-scale domains during neuronal maturation that align with topologically associating domains (TADs). These H3K27me3 “mega-domains” selectively encompass genes associated with alternative neural and non-neural lineages. While depletion of H3K27me3 in mature neurons has only modest effects on basal gene expression, it significantly increases neuronal activity-dependent c-FOS binding and induction of lineage-inappropriate genes within these mega-domains. Our findings reveal a previously unrecognized role for PRCs in establishing TAD-scale repressive chromatin domains during neuronal maturation, thereby safeguarding neuronal identity from external stimuli through broad silencing of alternative cell fate programs.

## Introduction

The generation of specialized cell types from pluripotent stem cells is driven by lineage-specific gene regulatory networks orchestrated by combinations of transcription factors (TFs)^1^. During development of the central nervous system, morphogen signaling and TF cascades establish neuronal identity with remarkable spatiotemporal precision^2,3^. Once specified, however, neuronal identity must be stably maintained throughout life. Mature neurons exhibit striking resistance to TF-mediated reprogramming and preserve their core transcriptional identity despite continuous exposure to environmental and activity-dependent stimuli^4,5^. These observations suggest that terminally differentiated cells are protected by epigenetic mechanisms that reinforce lineage identity and suppress alternative cell fate programs. However, the molecular basis of this stability remains poorly understood.

Polycomb group (PcG) proteins are evolutionarily conserved transcriptional repressors that play a central role in developmental gene regulation^6–8^. PcG proteins primarily function through two Polycomb repressive complexes (PRCs), PRC1 and PRC2, which deposit the repressive histone modifications H2AK119ub1 and H3K27me3, respectively^9–13^, and promote higher-order chromatin compaction^14–16^. PRCs are typically recruited to unmethylated CpG island-rich promoters^17–19^, where they spread via feed-forward recruitment of PRCs by recognition of their catalytic products^20–23^. In embryonic stem cells (ESCs) and progenitor populations, Polycomb-mediated repression is generally confined to localized chromatin domains spanning approximately 10 kb^23^ and is essential for controlling developmental transitions and lineage specification^24^.

In the mammalian central nervous system, PcG proteins are dynamically regulated during development. In early-stage neural progenitor cells (NPCs), PRCs maintain many neurodevelopmental genes in a poised state that permits subsequent activation. During lineage specification, PRCs regulate the temporal identity of NPCs by establishing stable repression at selected loci, thereby ensuring proper developmental progression^25–31^. Although recent studies have uncovered important functions of PcG proteins in postmitotic neurons^32–36^, the lack of longitudinal profiling during neuronal maturation has left a significant gap in our understanding. In particular, whether and how Polycomb-mediated chromatin regulation is reconfigured during neuronal maturation to stabilize long-term neuronal identity remains unknown.

Here, we show that Polycomb-mediated repression undergoes a profound reorganization during neuronal maturation. We find that H3K27me3 extends beyond promoter-proximal regions to form megabase-scale chromatin domains that coincide with topologically associating domains (TADs) and preferentially encompass genes associated with alternative neural and non-neural lineages. Functional perturbation of H3K27me3 reveals that these megabase-scale domains have only modest effects on basal transcription but play a critical role in restricting stimulus-dependent activation of lineage-inappropriate genes. Our findings reveal a previously unrecognized Polycomb repressive mode in mature neurons and suggest that TAD-scale H3K27me3 domains stabilize terminal cellular identity by broadly constraining alternative cell fate programs.

## Results

### H3K27me3 expands into megabase-scale domains during neuronal maturation

To investigate how Polycomb-mediated chromatin repression changes during neuronal maturation, we tracked a synchronized cohort of upper-layer excitatory neurons generated from E15 cortical progenitors by in utero electroporation (IUE) with a plasmid encoding SUN1-sfGFP (Fig. 1A). SUN1-sfGFP-expressing cells were isolated at four developmental stages (E15.5, E17, P1 and P7), spanning the transition from proliferative progenitors to mature postmitotic neurons (Fig. S1A). Quantitative normalization using spike-in controls revealed a progressive accumulation of H3K27me3 during neuronal maturation from E15 to P7 (Fig. 1B-1D). Strikingly, this increase was accompanied by a dramatic expansion of H3K27me3-enriched regions (Fig. 1E). Consistent with this global expansion, the size of individual H3K27me3-enriched regions increased markedly over the same period (Fig. 1F). Whereas H3K27me3 was relatively concentrated around promoter-proximal regions in neural progenitor cells, mature neurons displayed broader enrichment across gene bodies and intergenic regions (Fig. 1G and Fig. S1B).

**Figure 1.**
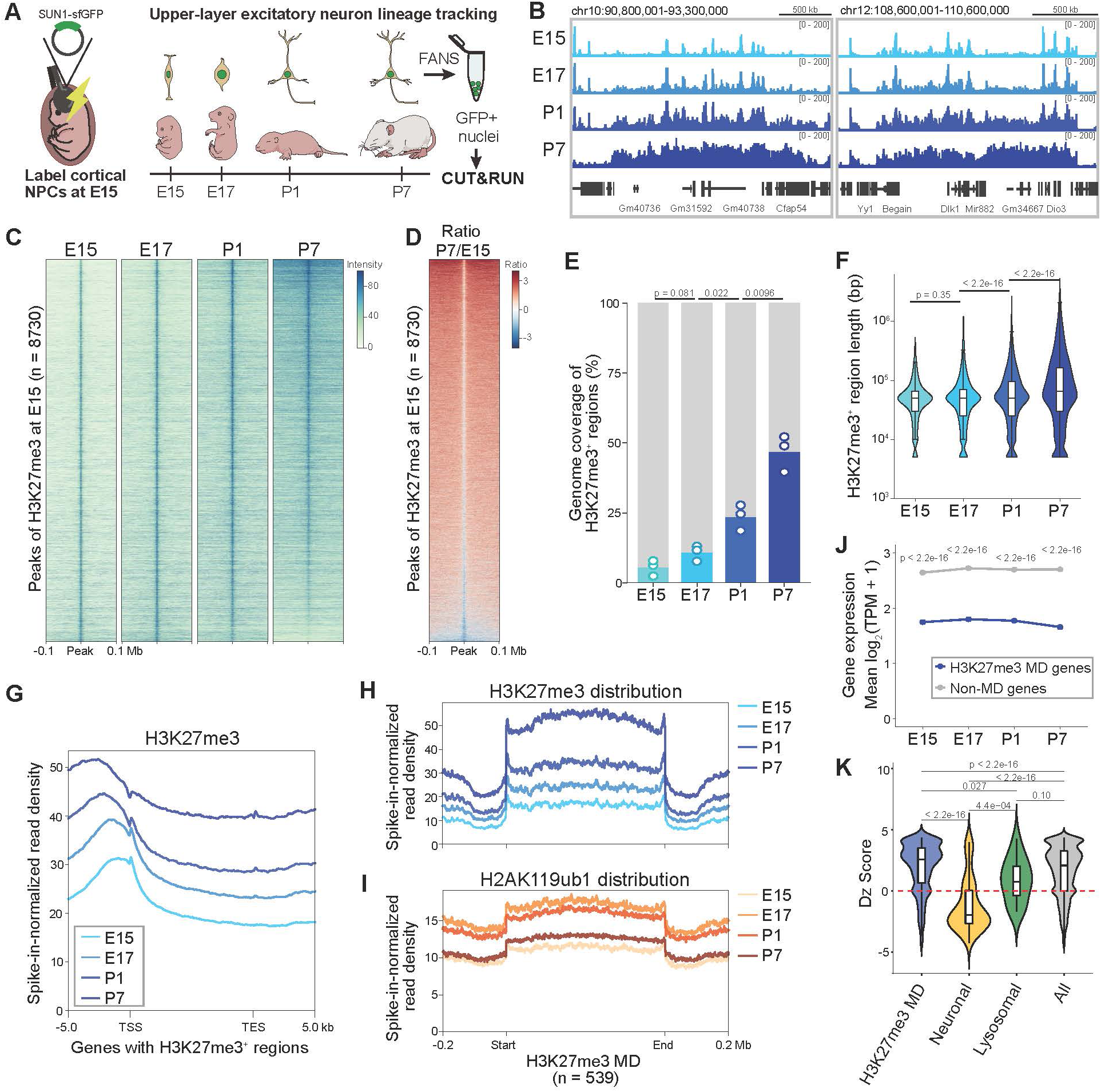
H3K27me3 increases to form mega-domains upon maturation of cortical neurons. (A) Schematic illustration of the nuclear labeling and isolation of the upper-layer cortical neuron lineage. In utero electroporation (IUE) was performed at embryonic day 15 (E15) to introduce a SUN1-sfGFP expression plasmid into the cortical ventricular zone, thereby labeling a subset of neural progenitor cells (NPCs) that generate upper-layer cortical neurons. Labeled nuclei were isolated at 0.5, 2, 5, and 11 days post-IUE, corresponding to E15.5, E17, P1, and P7 respectively. (B) Genome browser snapshots showing spike-in-normalized H3K27me3 signals at representative loci encompassing H3K27me3 target genes in SUN1-sfGFP^+^ nuclei across four developmental stages: E15.5, E17, P1, and P7 (sum of n = 3 biological replicates per stage). (C) Heatmap showing spike-in normalized H3K27me3 signals at H3K27me3 peaks defined at E15.5 SUN1-sfGFP^+^ labeled nuclei (n = 3 biological replicates per stage, 8730 regions). Rows are ordered by H3K27me3 signal at P7 in descending order. (D) Heatmap showing the log_2_-transformed P7/E15 ratio of H3K27me3 levels. Rows are ordered as in Figure 1C. (E) Bar graph showing the genomic coverage of H3K27me3^+^ regions (those within the top 20% of signal intensity across all developmental stages) at each developmental stage. H3K27me3^+^ regions were defined as genomic bins within the top 20% of signal intensity across all developmental stages. (F) Violin plot showing the length distribution of H3K27me3^+^ regions defined in Figure 1E at each developmental stage. (G) Metaplot showing the average spike-in-normalized H3K27me3 signal across H3K27me3-enriched genes, defined as the union of genes with H3K27me3^+^ regions identified at each developmental stage, and their 5-kb flanking regions. Signals are aligned at the TSS and TES. (H, I) Metaplots showing the average spike-in-normalized signal profiles of H3K27me3 (H) and H2AK119ub1 (I) across H3K27me3 mega-domains defined at P7 (539 regions), including flanking regions. Signals are aligned at the start and end of each domain. (J) Line plot showing the mean log_2_(TPM + 1) expression levels of H3K27me3 mega-domain (H3K27me3 MD) genes and non-MD genes at each developmental stage. (n = 4 biological replicates per stage, 50,000 nuclei per sample). (K) Violin plot showing Dz scores, which quantify the bias of gene expression toward non-brain versus brain tissues, for H3K27me3 mega-domain (MD) genes, neuronal genes, lysosome-related genes, and all genes. Lysosomal genes were used as a control gene set, as their expression shows little tissue specificity. Dz scores were calculated from juvenile mouse tissue CAGE profiles (P20–P30) in FANTOM5 as follows: Dz score = maximum Z-score among non-brain tissues − maximum Z-score among brain tissues. Positive and negative Dz scores indicate preferential expression in non-brain and brain tissues, respectively. Only genes represented in FANTOM5 were included. H3K27me3 MD genes, n = 3,803; neuronal genes annotated with “chemical synaptic transmission” (GO:0007268), n = 116; lysosome-related genes annotated with “regulation of lysosome organization” (GO:1905671), n = 14. Statistical significance was assessed by a Mann-Whitney U test followed by the Benjamini-Hochberg correction, and adjusted p-values are indicated.

We next examined whether H3K27me3 domains continue to evolve beyond the first postnatal week. Comparison of P7 and adult (P60) neurons revealed a further accumulation of H3K27me3 within a substantial fraction of H3K27me3 domains, together with the emergence of additional domains during postnatal maturation (Fig. S1C and S1D). A small subset of H3K27me3 domains, however, exhibited reduced H3K27me3 across gene bodies, accompanied by increased transcription of genes associated with neuronal functions between P7 and P60 (Fig. S1E-S1H). Together, these findings indicate that neuronal maturation is accompanied not only by increased H3K27me3 abundance but also by a large-scale reorganization of Polycomb-mediated chromatin repression.

To characterize these broad regions, we identified H3K27me3 domains exceeding 0.5 Mb in length in P7 neurons, hereafter termed H3K27me3 mega-domains. This analysis identified 539 mega-domains across the genome. H3K27me3 progressively accumulated within these domains during maturation (Fig. 1H), and the PRC1-associated modification H2AK119ub1 was similarly enriched (Fig. 1I), suggesting coordinated recruitment of the Polycomb repressive complexes PRC1 and PRC2. H3K27me3 mega-domains encompassed 6,058 genes whose expression remained substantially lower than the genomic average throughout development (Fig. 1J and S1I). To determine the tissue specificity of these genes, we defined the Differential Z (Dz) score as the difference between the maximum Z-score of gene expression across non-brain tissues and the maximum Z-score in brain tissue. Higher Dz scores indicate preferential expression in non-brain tissues, whereas lower Dz scores indicate brain-specific expression. Genes located within H3K27me3 mega-domains showed significantly higher Dz scores than the genomic background, indicating preferential expression in non-neural tissues (Fig. 1K). These findings suggest that H3K27me3 mega-domains preferentially target lineage-inappropriate gene programs rather than neuronal genes.

Collectively, these results identify H3K27me3 mega-domains as a prominent feature of mature neuronal chromatin that progressively expands during postnatal maturation while preferentially encompassing genes associated with alternative cellular identities.

### H3K27me3 mega-domains broadly emerge during tissue maturation and encompass lineage-inappropriate genes

The discovery of H3K27me3 mega-domains in mature cortical neurons prompted us to ask whether this chromatin reorganization is unique to the nervous system or reflects a general feature of tissue maturation. To address this question, we profiled H3K27me3 in fetal (E15) and adult (8–9 weeks) heart, renal cortex, liver, and lung using CUT&RUN. Whereas fetal tissues exhibited predominantly focal H3K27me3 peaks, adult tissues displayed broad H3K27me3 domains spanning large genomic regions (Fig. 2A). Quantification of genomic coverage confirmed a substantial increase in H3K27me3 occupancy in all adult tissues analyzed (Fig. 2B). The size of contiguous H3K27me3-enriched regions also increased during maturation (Fig. 2C). Thus, the transition from promoter-proximal Polycomb marking to large-scale H3K27me3 domains is not restricted to neurons but represents a widespread feature of postnatal tissue maturation.

**Figure 2.**
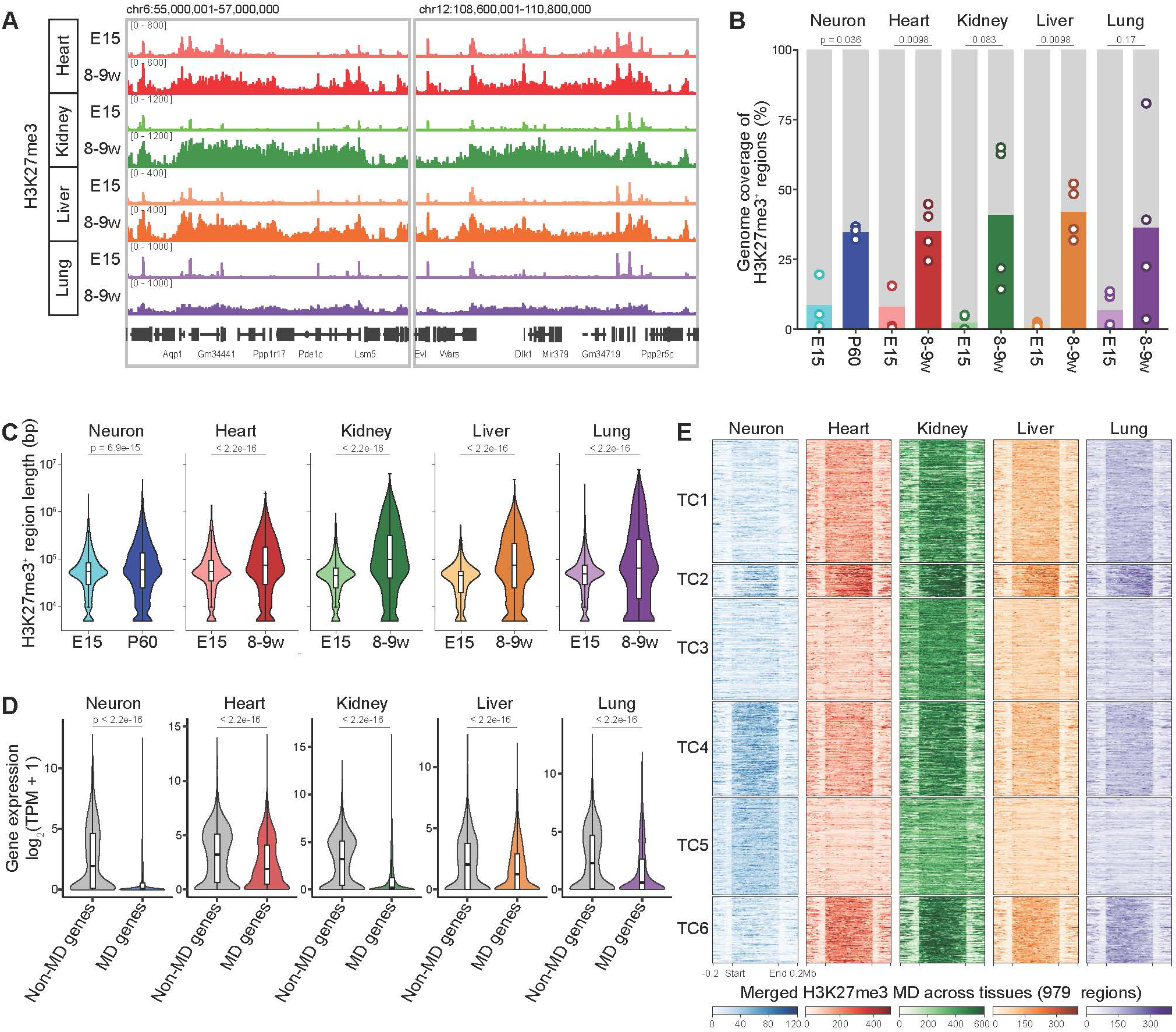
H3K27me3 increases in multiple adult tissues. (A) Genome browser snapshots of spike-in-normalized H3K27me3 signals in heart, kidney, liver and lung at E15.5 and 8-9 weeks of age at two representative loci. Signals are averaged across biological replicates (n = 4) for each tissue and stage. (B) Bar graph showing the genomic coverage of H3K27me3^+^ regions at each tissue and stage. H3K27me3^+^ regions were defined within each tissue as genomic bins within the top 20% of signal intensity across E15.5 and 8-9 weeks. Dots indicate individual biological replicates (n = 4). (C) Violin plots showing the length distribution of H3K27me3^+^ regions at E15 and the mature stage in neurons, heart, kidney, liver, and lung. H3K27me3^+^ regions were defined as in Figure 1E for neurons and as in Figure 2B for heart, kidney, liver, and lung. Numbers of regions analyzed at E15 and the mature stage, respectively, were as follows: neurons, 6,495 and 7,748; heart, 5,165 and 7,867; kidney, 2,415 and 6,071; liver, 1,479 and 8,675; lung, 4,933 and 6,212. (D) Violin plots showing gene expression levels of H3K27me3 mega-domain (MD) genes and all other genes (non-MD genes) in each tissue and developmental stage. Numbers of MD genes analyzed were as follows: neurons, 2,298; heart, 2,824; kidney, 3,392; liver, 3,005; lung, 2,689. (n = 2 biological replicates, 10,000 nuclei per sample for peripheral tissues. Data for neurons are the same as in Fig. S1E). (E) Heatmap showing spike-in-normalized H3K27me3 signals at mature-stage H3K27me3 mega-domains (MDs) across five tissues: neurons at P60 and heart, kidney, liver, and lung at 8–9 weeks of age. MDs were defined as the union of H3K27me3 MDs identified across the five tissues at the mature stage (n = 979 merged regions). Signals are shown across scaled MD bodies and ±0.2-Mb flanking regions. Rows were grouped by k-means clustering (k = 6) based on per-tissue mean signals and sorted within each cluster by decreasing mean signal. Statistical significance was assessed by a Mann-Whitney U test followed by the Benjamini-Hochberg correction.

We identified H3K27me3 mega-domains in each tissue to investigate the genes associated with these domains. RNA-seq analysis revealed that genes located within mega-domains (MD genes) were consistently expressed at lower levels than genes outside mega-domains (non-MD genes) across all tissues examined (Fig. 2D), indicating that these domains are associated with stable transcriptional repression. We next asked whether the genomic targets of H3K27me3 mega-domains are shared across tissues or tissue-specific. Clustering of H3K27me3 signals across merged H3K27me3 mega-domains identified both constitutive and tissue-selective patterns of domain occupancy (Fig. 2E). Several clusters displayed strong H3K27me3 enrichment across all tissues and were enriched for canonical developmental regulators, including Hox genes and keratin gene clusters (Fig. S2). In contrast, other clusters showed tissue-specific depletion of H3K27me3. Mega-domains enriched in non-neural tissues (TC3) were associated with synaptic genes, whereas domains not enriched in non-neural tissues (TC5) contained immune-related genes, likely reflecting the preservation of tissue-resident immune programs. These observations suggest a common organizational principle across tissues: H3K27me3 mega-domains preferentially encompass genes associated with alternative lineage programs while sparing genes required for the identity and function of the host tissue. Thus, large-scale Polycomb domain formation appears to be a general feature of tissue maturation that reinforces cellular identity through broad repression of lineage-inappropriate gene networks.

### H3K27me3 mega-domains align with pre-existing topologically associating domains

The sharp boundaries of H3K27me3 mega-domains across stages (Fig. 1G) prompted us to investigate whether their formation is constrained by higher-order chromatin architecture. To this end, we performed in situ Hi-C on E15 PAX6^+^NeuN^−^ neural progenitor cells (NPCs) and P7 NeuN^+^ neurons, and compared the resulting topologically associating domains (TADs) with H3K27me3 distribution. Remarkably, the boundaries of many H3K27me3 mega-domains coincided with TAD boundaries, as defined by peaks of the insulation score, in mature neurons (Fig. 3A and 3B). Moreover, most of these TAD boundaries were already present in E15 NPCs prior to mega-domain formation, indicating that the underlying chromatin architecture is established before neuronal differentiation and subsequently serves as a framework for H3K27me3 expansion during maturation. These observations suggest that H3K27me3 does not spread indiscriminately across the genome but accumulates preferentially within pre-existing topological domains.

**Figure 3.**
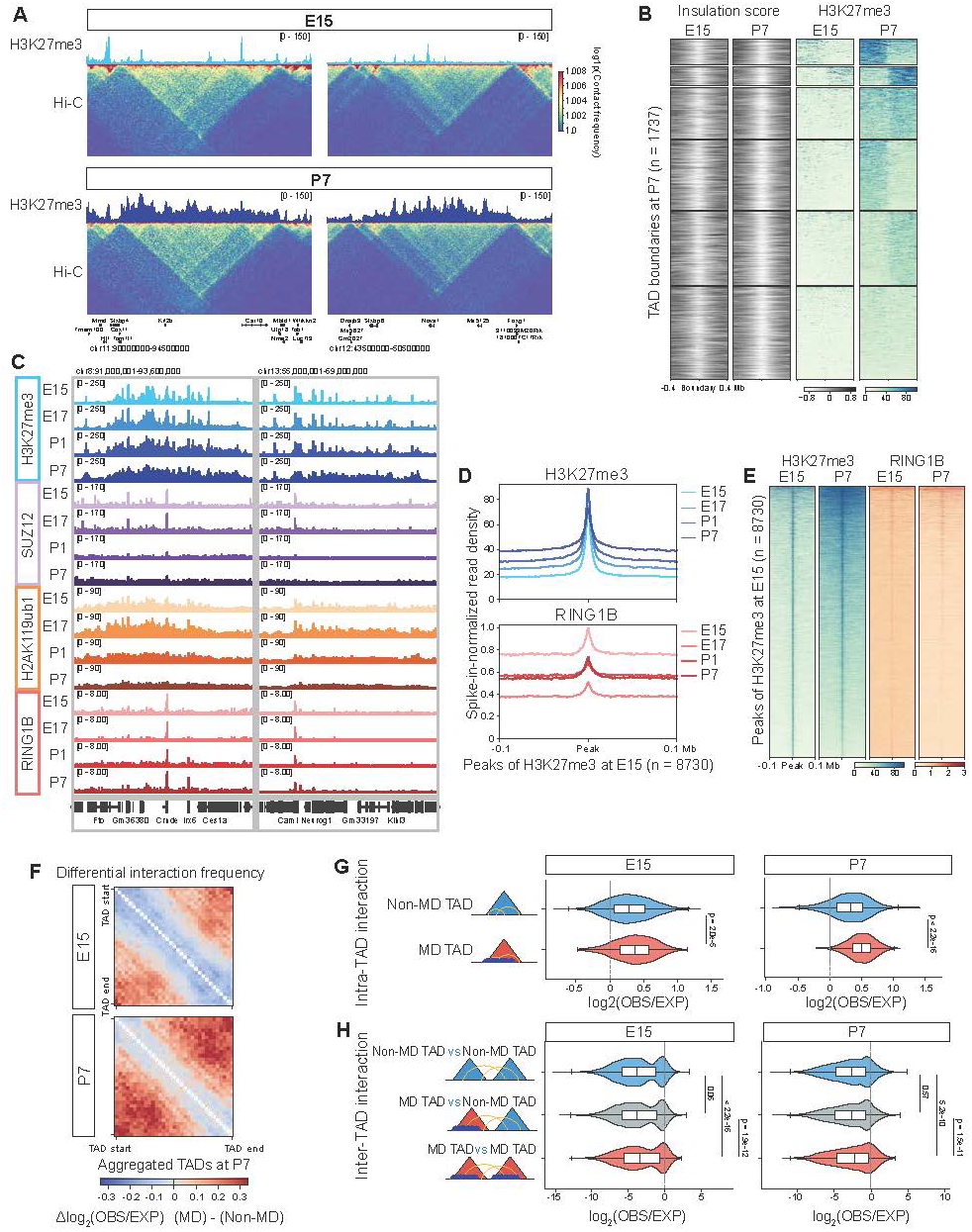
H3K27me3 mega-domains are associated with topologically associating domains. (A) Genome browser snapshots showing spike-in-normalized H3K27me3 distribution and Hi-C contact matrices at representative H3K27me3 mega-domain loci. Hi-C was performed using FANS-sorted E15 cortical NPCs (PAX6⁺/NeuN⁻) and P7 cortical neurons (NeuN⁺), whereas H3K27me3 CUT&RUN was performed using FANS-sorted SUN1-sfGFP⁺ nuclei from the in utero electroporated upper-layer cortical neuron lineage described in Figure 1A. Contact matrices are shown at 20-kb resolution after ICE normalization, with the color scale indicating log1p(contact frequency). Data were merged from n = 4 biological replicates per stage. (B) Heatmap showing the Hi-C insulation score and spike-in-normalized H3K27me3 signals at E15.5 and P7 TAD boundaries called at P7. Rows are centered on P7 TAD boundaries (n = 1,737) and show ±0.4-Mb flanking genomic regions. Rows were grouped by k-means clustering (k = 6). (C) Genome browser snapshots showing spike-in-normalized H3K27me3, SUZ12, H2AK119ub1 and RING1B signals at two representative regions in FANS-sorted SUN1-sfGFP⁺ cortical neuron lineage nuclei across four developmental stages: E15.5, E17, P1, and P7. Signals were averaged across n = 3 biological replicates per stage. Note that the H2AK119ub1 and SUZ12 signal was accompanied by a relatively high background, and SUZ12 intensity weakened as its expression declined during neuronal differentiation. (D) Metaplots showing the average spike-in-normalized signal profiles of H3K27me3 (top) and RING1B (bottom) signal profiles centered on H3K27me3 peaks identified in E15.5 SUN1-sfGFP⁺ nuclei across four developmental stages: E15.5, E17, P1, and P7 (n = 8,730 peaks; ±0.1-Mb flanking regions). (E) Heatmaps showing spike-in-normalized H3K27me3 and RING1B signals at H3K27me3 peaks identified in E15.5 SUN1-sfGFP⁺ nuclei (n = 8,730 peaks; ±0.1-Mb flanking regions).| (F) Heatmaps showing differential aggregate TAD analysis (ATA) between MD TADs and non-MD TADs at E15 and P7. MD TADs were defined as TADs overlapping P7 H3K27me3^+^ regions by ≥ 80% (n = 401), and non-MD TADs were defined as all other TADs (n = 1,279). The same classification was applied at both stages. Heatmaps show Δlog₂(observed/expected) contact frequency between MD and non-MD TADs, with red indicating stronger contacts in MD TADs. Only TAD bodies are shown, with the main diagonal masked. (G) Violin plots showing intra-TAD contact frequencies of MD TADs and non-MD TADs at E15 and P7, quantified as the mean log_2_(observed/expected) contact frequency across all valid bin pairs within each TAD; p-values were calculated using the Mann-Whitney U test. (H) Violin plots showing inter-TAD contact frequencies among MD and non-MD TAD pairs at E15 and P7. Inter-TAD contact frequency was quantified as the mean log_2_(observed/expected) contact frequency across all valid bin pairs between TADs on the same chromosome. P-values were calculated using the Mann-Whitney U test followed by Benjamini-Hochberg correction.

To determine whether this large-scale redistribution of H3K27me3 and H2AK119ub reflects altered Polycomb occupancy, we profiled the PRC2 component SUZ12 and the PRC1 component RING1B using CUT&RUN. Surprisingly, despite the dramatic expansion of H3K27me3 and H2AK119ub from discrete peaks in E15 progenitors into broad mega-domains in P7 neurons, SUZ12 and RING1B remained largely confined to their original peak regions even in P7 neurons (Fig. 3C-3E and S3A). The occupancy levels of SUZ12 and RING1B did not increased during neuronal maturation; instead, both decreased at many loci corresponding to H3K27me3 peaks identified in progenitors (Fig. 3C-3E and S3A). These findings indicate that mega-domain formation is not driven by widespread de novo recruitment of Polycomb complexes but rather reflects the propagation of Polycomb-mediated histone modifications from pre-existing Polycomb-bound sites across broader chromatin domains, potentially constrained by TAD architecture.

We next examined whether H3K27me3 accumulation within TADs is associated with chromatin interactions. In mature neurons, MD TADs (defined as TADs in which H3K27me3^+^ regions occupy more than 80% of the domain) exhibited significantly stronger intra-TAD interactions than non-MD TADs (Fig. 3F, 3G and S3B). Notably, this difference was much less pronounced in E15 progenitors, before extensive H3K27me3 spreading had occurred.

Among MD TADs, those with higher H3K27me3 levels showed stronger intra-TAD interactions, suggesting a positive association between H3K27me3 accumulation and intra-TAD interaction strength (Fig. S3C). Furthermore, H3K27me3-positive TADs interacted preferentially with one another, exhibiting elevated inter-TAD contact frequencies relative to interactions involving non-MD TADs (Fig. 3H). Together, these findings suggest that H3K27me3 mega-domains emerge within a pre-existing network of interacting chromatin domains and become increasingly concentrated within highly connected regions of three-dimensional nuclear space.

### Active transcription demarcates H3K27me3 mega-domain boundaries

Having found that H3K27me3 mega-domains are constrained by pre-existing TAD architecture, we next sought to identify the features that define their boundaries. The chromatin insulator CTCF was enriched at TAD boundaries, suggesting that boundary insulation may restrict the spreading of H3K27me3 (Fig. 4A). Given that transcriptional activity has been implicated in chromatin insulation^37^, we examined the distribution of active chromatin marks around H3K27me3 mega-domain edges. CUT&Tag profiling revealed a strong depletion of RNA Polymerase II occupancy within H3K27me3 mega-domains and a corresponding enrichment at their boundaries (Fig. 4B and 4C). Similarly, H3K4me3 was highly concentrated at domain edges, indicating the presence of active promoters flanking H3K27me3 mega-domains (Fig. 4B and 4D). Genes marked by H3K4me3 at the edges of H3K27me3 mega-domains were predominantly oriented away from the domains, suggesting that transcription is directed outward from the mega-domain interior (Fig. 4E-4G, S4A). Notably, many of these H3K4me3 peaks were already present in E15 neural progenitors before mega-domain formation, paralleling the developmental stability of TAD boundaries. Together, these findings identify actively transcribed loci as a defining feature of H3K27me3 mega-domain boundaries (Fig. 4H) and suggest that pre-existing transcriptionally active chromatin regions function as barriers that limit H3K27me3 propagation during neuronal maturation.

**Figure 4.**
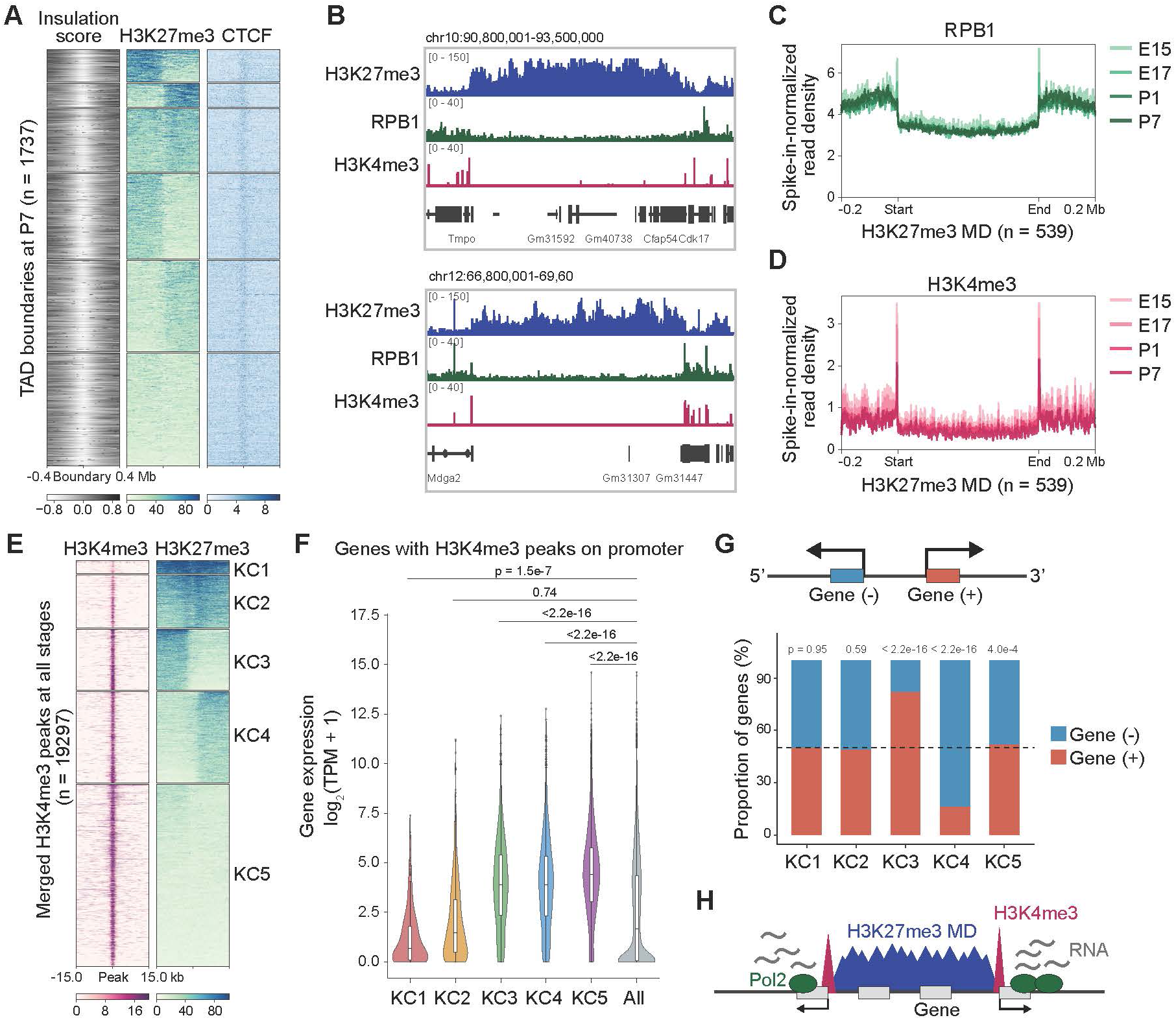
The boundaries of H3K27me3 mega-domains correlate with active transcription. (A) Heatmaps showing Hi-C insulation scores and spike-in-normalized H3K27me3 and CTCF signals around P7 TAD boundaries. Insulation scores and H3K27me3 signals are shown as in Figure 3B, together with spike-in-normalized CTCF signals at the same TAD-boundary clusters. Rows are centered on P7 TAD boundaries (n = 1,737) and show ±0.4-Mb flanking genomic regions. Rows were grouped by k-means clustering (k = 6) as in Figure 3B. (B) Genome browser snapshots showing spike-in-normalized H3K27me3, RPB1 and H3K4me3 signals at representative H3K27me3 MD loci at P7. (C, D) Metaplot showing average spike-in-normalized RPB1 (C) and H3K4me3 (D) signal profiles across P7-defined H3K27me3 MDs at four developmental stages: E15.5, E17, P1, and P7. Signals are aligned at the start and end of each H3K27me3 MD and shown with ±0.2-Mb flanking regions (n = 539 MDs). Profiles were averaged across n = 2 biological replicates per stage. (E) Heatmap showing spike-in-normalized H3K4me3 and H3K27me3 signals at P7 around merged H3K4me3 peaks identified across all four developmental stages (n = 19,297; ±15 kb flanks). Rows were grouped by k-means clustering (k = 5) based on P7 H3K4me3 and H3K27me3 signals. (F) Violin plot showing expression levels of genes associated with H3K4me3-marked promoters in each cluster defined in Figure 4E. Gene numbers per cluster were as follows: KC1, 266; KC2, 1,392; KC3, 2,387; KC4, 2,687; KC5, 6,163. Genes in each cluster were compared with all genes using a Mann-Whitney U test followed by the Benjamini-Hochberg correction; adjusted p-values are indicated. (G) Bar graph showing the proportion of plus- and minus-strand genes in each cluster defined in Figure 4E. Statistical significance was assessed by a binomial test followed by the Benjamini-Hochberg correction, and p-values are indicated. (H) Schematic model summarizing the relationship between H3K27me3 MDs, flanking H3K4me3 peaks, and active transcription.

### H3K27me3 mega-domains are embedded within a broader repressive chromatin environment

We next asked whether H3K27me3 mega-domains represent an extension of canonical Polycomb repression or instead occupy a distinct chromatin context in mature neurons. CUT&RUN profiling of Lamin B1 and H3K9me2 revealed increasing overlap between H3K27me3^+^ regions and lamina-associated domains (LADs) during neuronal maturation (Fig. 5A and 5B). In addition, Lamin B1 and H3K9me2 signals were significantly enriched within mega-domains relative to conventional H3K27me3 peaks (Fig. S5A and S5B). Overlap with H3K9me3-marked constitutive heterochromatin also increased during maturation, although to a lesser extent (Fig. S5C and S5D). We further investigated the distribution of non-CpG methylation (mCH), a repressive epigenetic modification that accumulates during neuronal maturation^38^, by analyzing whole-genome bisulfite sequencing (WGBS) data from cortical neurons at P40^38^. Notably, mCH accumulated substantially within H3K27me3 mega-domains in P7 neurons (Fig. 5C).

**Figure 5.**
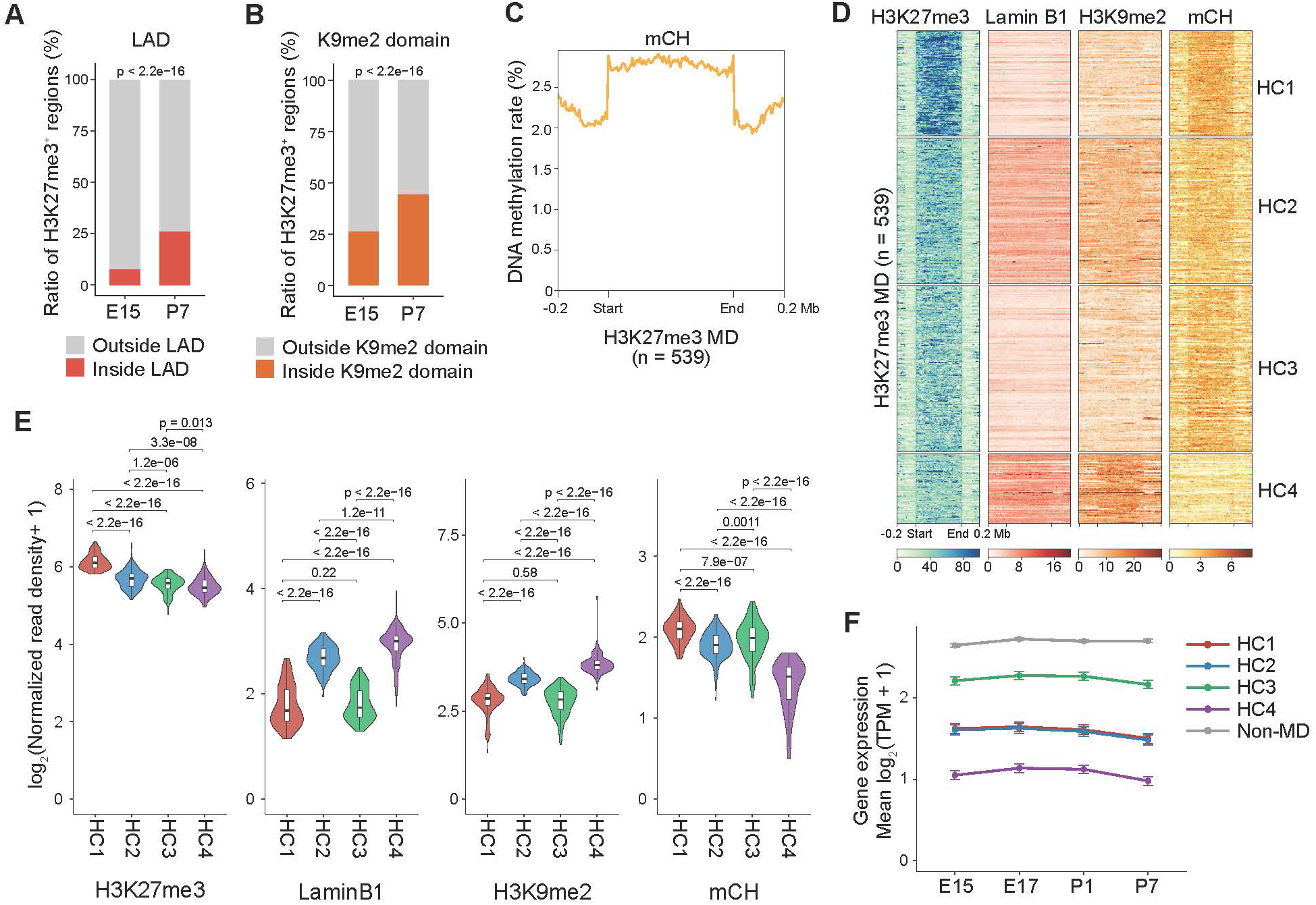
H3K27me3 mega-domains are associated with constitutive heterochromatin. (A, B) Bar graphs showing the proportions of H3K27me3^+^ regions located within Lamin B1-associated domains (LADs; A) or H3K9me2-enriched domains (K9me2 domains; B) at E15 and P7. LADs and K9me2 domains were defined from Lamin B1 and H3K9me2 profiles, respectively. P-values were calculated using Fisher’s exact test. n = 2 biological replicates per stage for Lamin B1 and H3K9me2. (C) Metaplot showing the average non-CpG methylation (mCH) rate in P40 cortical neurons from the frontal cortex (SRR921832) across P7-defined H3K27me3 MDs (n = 539). (D) Heatmaps showing H3K27me3, Lamin B1 and H3K9me2 signals and non-CpG methylation (mCH) rates across P7-defined H3K27me3 MDs. Rows were grouped by k-means clustering (k = 4) based on H3K27me3, Lamin B1, H3K9me2, and mCH levels within H3K27me3 MDs. (E) Violin plots showing the signal intensities of H3K27me3, Lamin B1, H3K9me2 and mCH levels in each cluster defined in Figure 5D. Statistical significance was assessed using a Mann-Whitney U test followed by the Benjamini-Hochberg correction; adjusted p-values are indicated. (F) Line plot showing mean expression levels of the genes in each H3K27me3 MD cluster defined in Figure 5D and non-MD genes across developmental stages. Data are shown as mean ± SEM across n = 4 biological replicates per stage. Gene numbers per cluster were as follows: HC1, 1,263 genes; HC2, 1,186 genes; HC3, 2,484 genes; HC4, 1,133 genes. Statistical significance was assessed by a Mann-Whitney U test followed by the Benjamini-Hochberg correction comparing non-MD genes to genes in each cluster, and p values are listed in the Supplemental Table.

To characterize the heterogeneity of mega-domains, we performed clustering based on H3K27me3, Lamin B1, H3K9me2, and mCH signals. Remarkably, most H3K27me3 mega-domains were categorized into LAD (Lamin B1 and H3K9me2)-enriched and/or mCH-enriched clusters (Fig. 5D and 5E). Genes contained within all mega-domain clusters were transcriptionally repressed relative to genes outside mega-domains (Fig. 5F). Notably, H3K27me3 mega-domains enriched only for Lamin B1 and H3K9me2 (HC4) were dominated by gene clusters, well-established targets of constitutive heterochromatin-mediated silencing. In contrast, H3K27me3 mega-domains enriched for mCH (HC1-HC3) were preferentially associated with genes involved in alternative developmental and tissue-specific programs, including skeletal, immune, renal and epithelial lineages (Fig. S5E).

Together, these findings indicate that H3K27me3 mega-domains do not simply represent enlarged versions of canonical Polycomb domains. Instead, they become integrated into a broader repressive chromatin landscape during neuronal maturation, where Polycomb repression cooperates with constitutive heterochromatin and neuronal DNA methylation to maintain stable silencing of lineage-inappropriate genes.

### H3K27me3 mega-domains safeguard neuronal identity against activity-induced transcriptional perturbation

To directly examine the function of H3K27me3 mega-domains, we acutely depleted H3K27me3 in postmitotic neurons after mega-domain establishment. To this end, we generated a doxycycline-inducible construct expressing the FLAG-tagged catalytic JmjC domain of the H3K27me3 demethylase KDM6B and introduced it into upper-layer cortical neurons by in utero electroporation (Fig. 6A and 6B). Induction at P7 for 24 h efficiently removed H3K27me3, as confirmed by immunostaining and CUT&RUN analyses (Fig. 6C-6F). RNA-seq following 24 h of JmjC overexpression identified only 133 upregulated genes and 7 downregulated genes (Fig. 6G), despite widespread depletion of a chromatin modification that occupies nearly half of the neuronal genome. Genes induced upon H3K27me3 removal were enriched for high baseline H3K27me3 levels (Fig. 6H), and a subset was located within H3K27me3 mega-domains (Fig. 6I). Analysis of tissue-wide expression patterns revealed that these genes were preferentially expressed in non-neural tissues and exhibited significantly higher Dz scores compared with the genomic background (Fig. 6J and 6K). These results indicate that H3K27me3 contributes to the repression of a limited set of lineage-inappropriate genes under basal conditions but is largely dispensable for maintaining transcriptional silencing across mega-domains.

**Figure 6.**
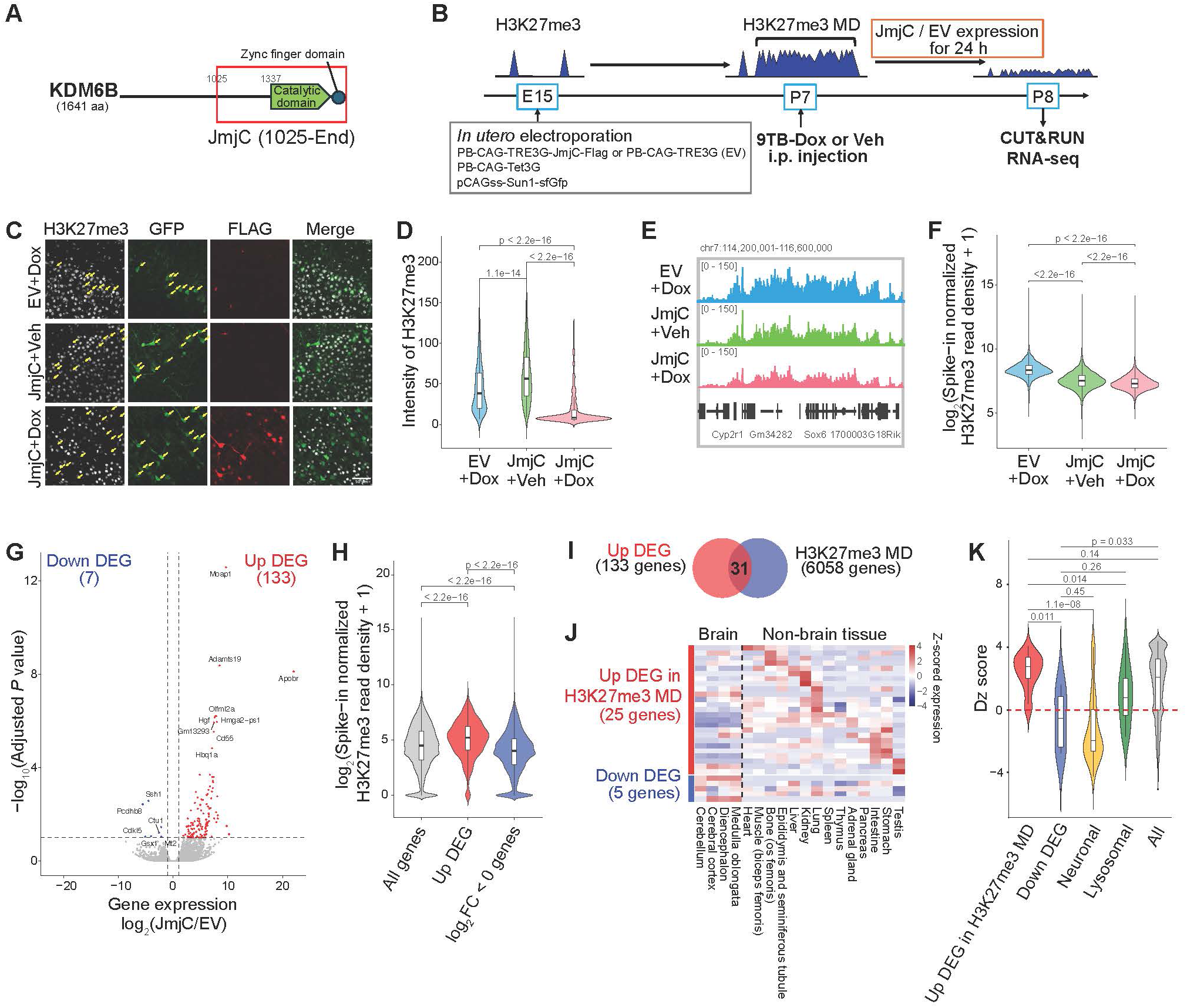
H3K27me3 depletion by KDM6B in neurons results in upregulation of non-neuronal genes. (A) Schematic illustration of the C-terminal JmjC fragment of mouse KDM6B used for overexpression in this study. The fragment spans amino acids 1,025 to 1,641 and contains the catalytic domain and zinc-finger domain. (B) Schematic illustration showing the experimental strategy for inducible JmjC expression. Plasmids encoding SUN1-sfGFP, Tet3G, and doxycycline-inducible JmjC-FLAG or control vector were introduced into cortical NPCs by IUE at E15.5. JmjC-FLAG expression was induced by 9TB-Dox injection at P7, and SUN1-sfGFP⁺ nuclei were isolated from the neocortex 24 h later at P8 for CUT&RUN and RNA-seq. (C) Representative immunofluorescence images showing H3K27me3 (grey), GFP (green), FLAG (red), and merged signals. Cells were co-transfected with GFP and either an empty vector (EV) or a doxycycline-inducible FLAG-tagged JmjC expression vector (JmjC), and treated with 9TB-Dox (Dox) or vehicle (Veh). Yellow arrows indicate GFP-positive transfected cells. Scale bar, 50 μm for all images. (D) Violin plot showing quantification of H3K27me3 fluorescence intensity in GFP-positive transfected cells. Numbers of cells analyzed from each brain were as follows: EV, n = 182, 130, and 129; JmjC + Veh, n = 70, 287, and 97; JmjC + Dox, n = 122, 23, and 50. (E) Genome browser snapshot showing the H3K27me3 signals in empty vector (EV) control samples (blue) and JmjC-expressing samples treated with vehicle (green) or 9TB-Dox (red). (F) Violin plot showing H3K27me3 read density at P8 across P7 H3K27me3^+^ regions defined in Figure 1D (n = 7,132 regions). Read counts were normalized by region length and spike-in ratio. (G) Volcano plot showing differential gene expression between JmjC-expressing samples (*n* = 6) and empty vector (EV) control samples (*n* = 4) at P8. Differentially expressed genes were defined as genes with p < 0.1 and an absolute fold change ≥ 2, and are highlighted. 10,000 nuclei per sample. (H) Violin plot showing H3K27me3 read density over gene bodies of all genes, upregulated DEGs in JmjC-expressing samples relative to empty vector (EV) control samples, and genes with log_2_ fold change < 0. Read density was normalized by gene length and spike-in ratio and is shown as log_2_(read density + 1). Statistical significance was assessed using a Mann–Whitney U test; p-values are indicated.| (I) Venn diagram showing the overlap between upregulated differentially expressed genes (upregulated DEGs) in JmjC-expressing samples relative to empty vector (EV) control samples and H3K27me3 MD genes. (J) Heatmap showing z-score-normalized expression of selected gene sets across juvenile mouse tissues (P20–P30) using FANTOM5 CAGE expression profiles. Gene sets include upregulated DEGs among H3K27me3 MD genes upon JmjC overexpression (25 genes) and downregulated DEGs upon JmjC overexpression (5 genes). Columns represent juvenile brain and non-brain tissues. (K) Violin plot showing the distribution of Dz scores for the indicated gene categories. For each gene, the Dz score was calculated as the maximum Z-score across non-brain tissues minus the maximum Z-score across brain tissues. Positive and negative Dz scores indicate preferential expression in non-brain and brain tissues, respectively. Neuronal genes annotated with “chemical synaptic transmission” (GO:0007268; 216 genes), and lysosomal genes annotated with “regulation of lysosome organization” (GO:1905671; 14 genes).

We next asked whether the function of H3K27me3 mega-domains becomes more apparent under conditions that challenge transcriptional homeostasis. To test this possibility, we induced neuronal activity with kainic acid (KA) following JmjC overexpression (Fig. 7A and S6A). Genome-wide analyses revealed markedly enhanced transcriptional derepression compared with unstimulated conditions (Fig. 6G, 7B and 7C). Notably, genes preferentially expressed outside the brain (Dz > 1) showed significantly greater derepression than brain-enriched genes (Fig. 7D). Among activity-responsive upregulated genes following H3K27me3 removal, 91 were located within H3K27me3 mega-domains (Fig. 7E), and these genes were strongly enriched for non-neural expression programs (Fig. S6C).

**Figure 7.**
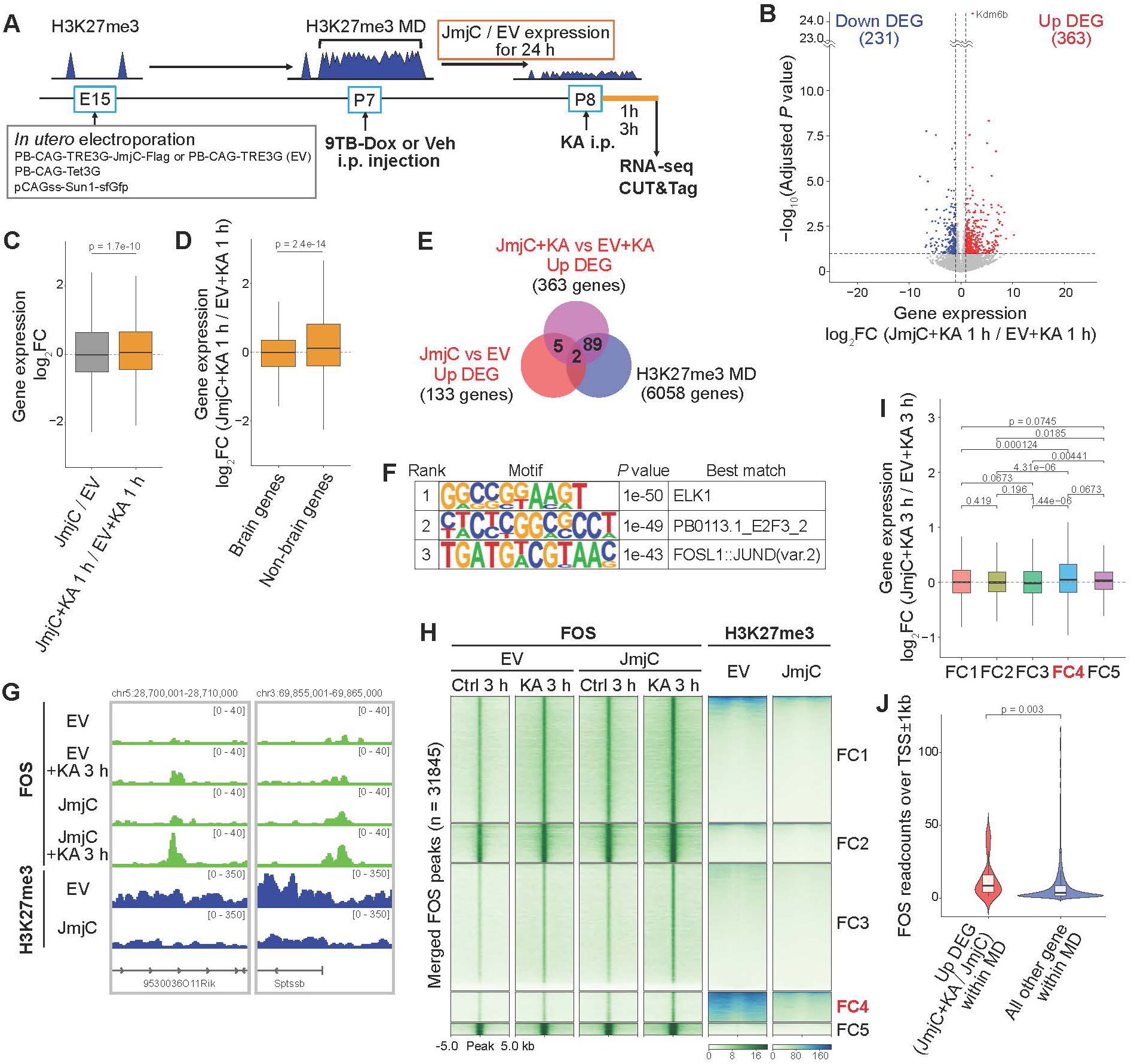
Activity-dependent transcription is dysregulated under H3K27me3 depletion. (A) Schematic illustration showing the experimental strategy for inducible JmjC expression followed by kainic acid (KA)-induced neuronal stimulation. Plasmids encoding SUN1-sfGFP, Tet3G, and doxycycline-inducible JmjC-FLAG or control vector were introduced into cortical NPCs by in utero electroporation (IUE) at E15.5. JmjC-FLAG expression was induced by intraperitoneal injection of 9TB-Dox at P7. 24 hrs later, mice were injected intraperitoneally with KA, and SUN1-sfGFP⁺ nuclei were isolated from the neocortex 1 h or 3 h after KA injection for downstream analyses. (B) Volcano plot showing differential gene expression between KA-injected JmjC-expressing samples (n = 6) and KA-injected empty vector (EV) control samples (n = 4) at P8. Differentially expressed genes were defined as genes with P < 0.1 and absolute fold change ≥ 2, and are highlighted. 10,000 nuclei per sample. (C) Box plot showing log_2_ fold changes in gene expression for all genes in two comparisons: JmjC-expressing samples versus empty vector (EV) control samples without KA injection, and KA-injected JmjC-expressing samples versus KA-injected EV control samples 1 h after KA injection. (D) Box plot showing log_2_ fold changes in gene expression for brain-biased genes and non-brain-biased genes, defined by Dz scores as described in Figure 1K, in KA-injected JmjC-expressing samples relative to KA-injected empty vector (EV) control samples 1 h after KA injection. Brain-biased and non-brain-biased genes were defined as genes with Dz < −1 (n = 2,272) and Dz > 1 (n = 12,398), respectively. (E) Venn diagram showing the overlap among upregulated DEGs identified in KA-injected JmjC-expressing samples relative to KA-injected empty vector (EV) control samples 1 h after KA injection, upregulated DEGs identified in JmjC-expressing samples relative to EV control samples, and H3K27me3 mega-domain (MD) genes. (F) Table showing the result of de novo motif analysis performed using HOMER on the promoter regions of upregulated DEGs in KA-injected JmjC-expressing samples relative to KA-injected control samples. Promoter regions were defined as −100 to +100 bp around the TSS. “Best match” indicates the most similar known motif in the HOMER database. (G) Genome browser snapshots showing FOS and H3K27me3 signals at a representative H3K27me3 mega-domain (MD) locus in empty vector (EV) control, KA-injected EV control, JmjC-expressing, and KA-injected JmjC-expressing samples. KA-injected samples were collected 3 h after KA injection. (H) Heatmaps showing spike-in-normalized FOS and H3K27me3 signals centered on merged FOS peaks (n = 31,845) in empty vector (EV) control and JmjC-expressing samples following KA injection. FOS signals were analyzed 3 h after KA injection. Rows show ±5 kb regions around FOS peak centers and were grouped into 5 clusters using two-step k-means clustering: an initial clustering (k=4) based on FOS signals, followed by sub-clustering of Cluster FC3+FC4 (k=2) based on both FOS and H3K27me3 signals across samples. (I) Box plot showing log_2_ fold changes in gene expression for genes associated with FOS peaks in each cluster defined in Figure 7H, comparing KA-injected JmjC-expressing samples with KA-injected EV control samples 3 h after KA injection. P-values were calculated using a Mann–Whitney U test followed by Benjamini–Hochberg correction. (n = 3 biological replicates per condition, 50,000 nuclei per sample). (J) Violin plot showing FOS read counts within ±1 kb of the TSS for upregulated DEGs within H3K27me3 mega-domains (MDs) and all other MD genes 3 h after KA injection. Upregulated DEGs were defined from the JmjC + KA versus JmjC comparison. P-value was calculated using a Mann–Whitney U test.

We then asked whether specific classes of mega-domains were particularly sensitive to H3K27me3 removal. Other-lineage genes residing in H3K27me3 mega-domain clusters HC1 and HC3, which are enriched for mCH but show limited association with Lamin B1 and H3K9me2, exhibited the strongest JmjC-dependent derepression following neuronal activation (Fig. S6E). These findings suggest that repression of many lineage-inappropriate genes remains directly dependent on H3K27me3 despite the presence of mCH-associated repressive mechanisms.

To identify transcription factors capable of engaging these newly accessible loci, we performed de novo motif analysis around the transcription start sites (TSSs) of genes upregulated by JmjC overexpression under KA stimulation. This analysis revealed significant enrichment of ELK1 and FOSL1::JUND motifs (Fig. 7F), both of which act downstream of activity-dependent signaling pathways. We therefore performed c-FOS CUT&Tag and indeed identified a c-FOS peak cluster (FC4) in which c-FOS occupancy was significantly increased 3 h after KA administration, but only when H3K27me3 had been eliminated by JmjC overexpression (Fig. 7G and 7H). The genes in FC4 were upregulated upon H3K27me3 depletion and were functionally associated with non-neuronal lineages (Fig. 7I and Fig. S6G). Moreover, c-FOS occupancy was greater at the promoters of genes upregulated by KA stimulation in the presence of JmjC overexpression, compared with other genes located within H3K27me3 mega-domains (Fig. 7J). Together, these results suggest that removal of H3K27me3 renders a subset of previously inaccessible lineage-inappropriate genes responsive to endogenous activity-induced transcription factors, including c-FOS.

## Discussion

The present study uncovers a previously unrecognized mode of Polycomb organization that emerges during terminal neuronal maturation in vivo. By selectively isolating upper-layer cortical neurons throughout development, we identified the progressive formation of hundreds of TAD-scale, megabase-sized H3K27me3 mega-domains that are largely absent in neural progenitor cells. Such large-scale reorganization of Polycomb domains has not been observed in previous studies of neuronal differentiation using pluripotent stem cell-derived in vitro systems^39^, suggesting that this chromatin transition depends on developmental features that are not fully recapitulated in culture. One possibility is that tissue-specific microenvironmental cues present in vivo are required for mega-domain formation. Importantly, comparable mega-domains were observed across all mature tissues examined, suggesting that this phenomenon is not unique to neurons but may represent a general feature of terminal tissue maturation.

A central conceptual advance of this study is that H3K27me3 mega-domains function not primarily as constitutive repressors, but as stimulus-responsive epigenetic safeguards that preserve neuronal identity. Acute depletion of H3K27me3 by JmjC overexpression induced only a limited number of genes, and those located within H3K27me3 mega-domains were preferentially associated with non-neural lineages. These findings are consistent with previous reports showing that global reduction of H3K27me3 following Ezh1/2 deletion in neurons causes surprisingly modest transcriptional derepression^32,40^. However, while H3K27me3 depletion alone produced only modest transcriptional changes, its functional importance became much more evident following neuronal activation.

Activity-induced transcription in the absence of H3K27me3 resulted in widespread derepression of lineage-inappropriate genes. The activity-dependent transcription factor c-FOS accumulated at these derepressed genes upon H3K27me3 depletion, indicating that endogenous activity-dependent transcription factors gain access to otherwise repressed loci when H3K27me3 is removed. These observations suggest that H3K27me3 mega-domains do not primarily function as constitutive repressors under basal conditions. Instead, H3K27me3 mega-domains appear to buffer physiological transcriptional responses, preventing endogenous activity-dependent transcription factors from aberrantly engaging alternative lineage programs. Because AP-1 and ELK1 are activated not only by neuronal activity but also by diverse extracellular signals, including neurotrophic factors, neurotransmitters, and inflammatory stimuli^41^, H3K27me3 mega-domains may provide a broader protective mechanism that prevents inappropriate activation of alternative lineage programs in response to fluctuating environmental inputs.

Importantly, a large subset of genes contained within H3K27me3 mega-domains still remained transcriptionally silent even after acute H3K27me3 depletion. This observation suggests that additional repressive mechanisms cooperate with the Polycomb system to maintain stable silencing. Consistent with this idea, we found that H3K27me3 mega-domains frequently overlap with lamina-associated domains, H3K9 methylation, and neuronal non-CpG DNA methylation. Genes located within Lamin B1-enriched mega-domains were particularly resistant to derepression following H3K27me3 removal, in agreement with previous studies demonstrating redundant functions of H3K9me2, H3K9me3, and H3K27me3 in heterochromatin-mediated silencing^42–44^. Thus, H3K27me3 mega-domains appear to function as components of a multilayered repressive chromatin environment rather than as isolated Polycomb structures.

Another unexpected feature of mature H3K27me3 mega-domains is their extensive coexistence with mCH, H3K9 methylation, and lamina-associated chromatin. DNA methylation and H3K27me3 have traditionally been viewed as mutually exclusive chromatin features, especially at CpG island promoters where DNMT3-mediated methylation antagonizes PRC2 recruitment^38,45–47^. However, recent studies have shown that H3K27me3 and DNA methylation can coexist outside CpG islands. In neurons, mCH and hydroxymethylation accumulate extensively during postnatal maturation and are enriched within specific topological domains^38,48^. Our findings raise the possibility that mCH and H3K27me3 cooperate to establish repressive TAD-scale chromatin environments in mature neurons. Whether these pathways act redundantly, synergistically, or hierarchically during mega-domain formation and maintenance remains an important question for future investigation.

Previous work has primarily viewed PcG proteins as developmental regulators whose repressive activity must be relieved to permit neuronal maturation^34,49^. Our findings instead reveal a complementary architectural role for PcG proteins after differentiation, in which TAD-scale H3K27me3 mega-domains contribute to the long-term preservation of mature neuronal identity. This maintenance function likely requires a more stable and durable mode of repression than the transient gene silencing classically associated with developmental transitions. H3K27me3 mega-domains may provide such stability through multiple mechanisms. Beyond the accumulation of Polycomb-mediated histone modifications across large genomic regions, these domains frequently colocalize with additional repressive chromatin features as mentioned above, and exhibit enhanced intra-TAD interactions. Consistent with this interpretation, Polycomb bodies visualized by RING1B immunostaining appeared more discrete and condensed in mature neurons than in neural progenitors (Fig. S6H). It is therefore possible that maturation-associated changes in Polycomb composition or cooperation with other repressive factors promote higher-order chromatin compaction. Notably, several Polycomb components implicated in chromatin condensation, including members of the CBX and PHC families, are upregulated during neuronal differentiation^50^ and may contribute to the establishment of these highly stable repressive structures.

Our data also challenge the prevailing model for Polycomb domain formation. Based on studies in embryonic stem cells, one might predict that progressive recruitment of PRC1 and PRC2 through canonical feed-forward interactions would drive domain expansion^20–22^. However, our data do not support this model, given that H3K27me3 accumulation occurred uniformly across entire TADs rather than spreading progressively from nucleation sites, and that the catalytic Polycomb complexes themselves remained confined to narrow peaks even after mega-domains had formed. Instead, we propose that Polycomb complexes anchored at discrete nucleation sites repeatedly modify chromatin throughout the surrounding TAD through frequent three-dimensional chromatin contacts (see Graphical Abstract). In this framework, pre-existing TAD architecture provides the structural template that constrains and facilitates domain-wide accumulation of H2AK119ub1 and H3K27me3.

An important unresolved question concerns how specific TADs are selected for mega-domain formation. Our analyses suggest that actively transcribed loci positioned near TAD boundaries can function as barriers to H3K27me3 accumulation. In addition, other chromatin features, including mCH and lamina association, may contribute to defining TADs that are permissive for large-scale Polycomb accumulation. Elucidating how these architectural and epigenetic features interact to determine mega-domain identity will be an important direction for future studies.

In conclusion, our study uncovers a previously unrecognized mode of Polycomb function in terminally differentiated cells. During maturation, Polycomb-mediated repression expands into TAD-scale H3K27me3 mega-domains that broadly silence lineage-inappropriate genes and safeguard cellular identity against transcriptional perturbation. Our findings therefore identify H3K27me3 mega-domains as a conserved epigenetic architecture for stable cell identity maintenance and suggest that controlled modulation of these structures may help overcome epigenetic barriers to cellular reprogramming, thereby facilitating regenerative medicine approaches.

## Author contribution

Conceptualization, N.Y., C.I., K.O., H.S. and Y.G.; Investigation, N.Y., C.I., K.H., K.O. and H.S.; Formal Analysis, N.Y., C.I., and H.S.; Data Curation, N.Y. and H.S.; Writing – Original Draft, N.Y., H.S., and Y.G.; Writing – Review & Editing, all authors; Visualization, N.Y., C.I., H.S.; Supervision, H.S. and Y.G.; Project Administration, H.S. and Y.G.; Funding Acquisition, N.Y., K.O., H.S. and Y.G.

## Acknowledgements

We thank H. Koseki (RIKEN) for providing protein A/G-MNase; and members of the Gotoh laboratory for discussion. This work was supported by JSPS KAKENHI (Grant Numbers JP22H00431, JP25H00435, JP24H02322, JP22KK0107, JP25K22503, and JP25K24466 to Y.G.; JP25K18471 and JP22K15118 to H.S.; JP24K09656 to K.O. and JP24KJ0951 to N.Y.); by AMED (Grant Numbers JP24gm1310004 [AMED-CREST] and JP25wm0625223 to Y.G.; and JP23wm0525035 and JP223fa627001 to H.S.); by the Secom Science and Technology Foundation and the Uehara Memorial Foundation (to H.S.); by the World Premier International Research Center Initiative (WPI), MEXT, Japan, through the International Research Center for Neurointelligence (WPI-IRCN); by Rikaken Holdings Co., Ltd. (to N.Y.); and by the International Graduate Program of Innovation for Intelligent World (IIW).

## STAR★Methods

### Key resources table

->Attached Excel file

## RESOURCE AVAILABILITY

### Lead contact

Further information and requests for resources and reagents should be directed to and will be fulfilled by the lead contact, Hiroki Sugishita and Yukiko Gotoh.

### Materials availability

Commercially available reagents are listed in the key resources table. All plasmids generated in this study are available on request.

### Data and code availability

The original data and source codes reported in this paper are available from the lead contact upon request.

### Experimental model and study participant details Mice

All animal experiments were approved by the Animal Care and Use Committee of the Graduate School of Pharmaceutical Sciences, the University of Tokyo (approval numbers: P25-8 and P30-4), and were performed in accordance with the University of Tokyo guidelines for the care and use of laboratory animals and the ARRIVE guidelines. Slc:ICR (ICR) mice were obtained from Japan SLC, Inc. (Shizuoka, Japan). Mice were housed in a climate-controlled facility maintained at 23 ± 3 °C and 50 ± 15% relative humidity under a 12-h light/dark cycle, with food and water provided ad libitum. The day of vaginal plug detection was designated as embryonic day 0.5 (E0.5), and the day of birth was designated as postnatal day 0 (P0). Both male and female mice were used in this study, and sex was not considered as a biological variable. For the analysis of bulk tissues, only male mice were used.

### Plasmid construction

To generate the inducible JmjC expression construct encoding the N-terminally Flag- and SV40 NLS-tagged catalytic domain of mouse Kdm6b, a multi-step cloning strategy was employed. First, a cDNA fragment encompassing the C-terminal 1,025 amino acids of Kdm6b (corresponding to amino acids E–R) was amplified by PCR from Addgene plasmid #100278 using the forward primer 5’-atgcgaattcgccaccatggactacaaagacgatgacgacaag-3’ and the reverse primer 5’-gcatgcggccgctcatcgagacgt-3’. To introduce the SV40 NLS sequence, a second PCR was performed using the first PCR product as a template with the forward primer 5’-acgatgacgacaagggatccccaaagaagaagcggaaggtcggtatccacggagtcc-3’ and the reverse primer 5’-cctggatctcggatccggctgctgggactccgtggataccgaccttcc-3’. The first PCR product was digested with BamHI and subsequently assembled with the second PCR product via In-Fusion assembly. Finally, the resulting construct was digested with EcoRI and NotI, and ligated into the corresponding restriction sites of the PB-TRE3G PiggyBac vector^51^ to yield PB-TRE3G-JmjC.

To generate the pCAG-SUN1(trunc.)-sfGFP plasmid for cell-type-specific nuclear labeling, an N-terminally truncated variant of the mouse Sun1 cDNA, as previously described^52^, was fused in-frame to the N-terminus of superfolder GFP (sfGFP) and inserted into the pCAGGS vector^53^ under the control of the CAG promoter.

## METHOD DETAILS

### In utero electroporation (IUE)

In utero electroporation was performed as previously described^54^ with minor modifications. Timed-pregnant ICR mice at embryonic day 15 (E15) were anesthetized by intraperitoneal injection of a mixed anesthetic solution containing medetomidine (Domitor, Nippon Zenyaku Kogyo), midazolam (Sandoz), and butorphanol (Vetorphale, Meiji Seika Pharma, Cat# 28-0192). Plasmid DNA in PBS containing 0.01% Fast Green (CAY, Cat# 14335) was injected into the lateral ventricle of each embryo through a pulled glass capillary. Electric pulses were delivered using 5-mm forceps-type platinum electrodes (CUY650P5, Nepa Gene) and a NEPA21 Type II electroporator (Nepa Gene), consisting of three poring pulses (40 V, 30-ms duration, 50-ms interval, 10% decay rate) followed by four transfer pulses (10 V, 30-ms duration, 50-ms interval, 40% decay rate). After surgery, anesthesia was reversed by subcutaneous injection of atipamezole (Antisedan, Nippon Zenyaku Kogyo).

For nuclear labeling of upper-layer excitatory neurons across developmental stages, a mixture of pCAGEN-Sun1(trunc)-sfGFP (1 μg/μL) and pCAGEN-mCherry (1 μg/μL) was electroporated at E15.25, and cortices were dissected at E15.75 (12 h post-IUE), E17, P1, or P7.

For doxycycline-inducible overexpression of the Kdm6b catalytic domain (JmjC), a mixture of PB-CAG-TRE3G-JmjC (1.5 μg/μL), PB-CAG-Tet3G (1.2 μg/μL), pCAGEN-Sun1(trunc)-sfGFP (0.8 μg/μL), and pCAGEN-mCherry (0.5 μg/μL) was electroporated at E15. To induce JmjC expression, 9-t-butyl-doxycycline (9TB-dox, 20 mg/kg) was administered intraperitoneally to pups at P7, and cortices were dissected 24 h later. PB-CAG-TRE3G empty vector was co-electroporated in place of PB-CAG-TRE3G-JmjC as a control. For neuronal activation experiments, kainic acid (KA; Sigma-Aldrich, Cat# K0250; 0.4 mg/mL in D-PBS) was administered intraperitoneally at 4 mg/kg at P8 (24 h after the initial 9TB-dox injection), and cortices were dissected 1 h or 3 h after KA administration.

### Immunohistofluorescence analysis

For immunohistochemical staining of electroporated brain sections, mice were transcardially perfused with ice-cold 4% paraformaldehyde (Wako, Cat# 162-16065) in PBS. The brain was then removed, exposed to the same fixative for 1.5 h at 4 °C, equilibrated with 30% sucrose in PBS, embedded in O.C.T. compound (Tissue TEK, Cat# 4583), and frozen. Coronal cryosections (thickness of 12 μm) were subjected to antigen retrieval with Target Retrieval Solution (Agilent, Cat# S1699) at 105 °C for 10 min, and were exposed to Tris-buffered saline containing 0.1% Triton X-100 and 3% BSA (blocking buffer) for 1 h at room temperature, incubated first overnight at 4 °C with primary antibodies in blocking buffer and then for 1 h at room temperature with Alexa Fluor-conjugated secondary antibodies (1:1,000 dilution, Thermo Fisher) and Hoechst 33342 (1:1,000 dilution, Molecular Probes) in blocking buffer, and mounted in Mowiol (Calbiochem). Fluorescence images were obtained with a laser confocal microscope (Zeiss LSM 880) and were processed with the use of ZEN (Zeiss), and ImageJ (NIH) software. Antibodies are listed in the key resources table. For the quantification of H3K27me3 and RING1B fluorescence, regions of interest (ROIs) were segmented using Cellpose, and the fluorescence intensity was measured using ImageJ. Quantification used numpy, scipy, and scikit-image.

### Nuclear extraction and FANS

Two nuclear isolation protocols were used depending on the downstream application: a Dounce homogenization protocol for Hi-C and CUT&RUN of peripheral tissues, and a Percoll gradient-based protocol for CUT&RUN and RNA-seq of neocortical tissue. For Dounce homogenization, snap-frozen tissues were homogenized in a 1-mL Dounce homogenizer in ice-cold Nuclear Extraction Buffer (10 mM Tris-HCl pH 7.6, 0.25 M sucrose, 25 mM KCl, 5 mM MgCl₂, 0.1% Triton X-100) supplemented with 1 mM DTT, 1× protease inhibitor cocktail (Roche), and 0.4 U/μL RNase inhibitor (TOYOBO). Nuclei were pelleted at 500 × g, washed in 1% BSA/PBS containing the same inhibitors, and filtered through a 35-μm cell strainer (Falcon). For the Percoll gradient, snap-frozen neocortices were homogenized in 54% Percoll (Wako, Cat# 592-09051) in Isolation Buffer (0.25 M sucrose, 250 mM KCl, 50 mM MgCl₂, 125 mM Tris-HCl pH 7.4) containing 0.1% NP-40, sequentially underlaid with 31% and 35% Percoll in Isolation Buffer, and centrifuged at 20,000 × g for 10 min at 4°C in a swinging-bucket rotor. The interphase nuclear fraction was collected, washed twice in 0.1% BSA/PBS, and filtered through a 35-μm cell strainer (Falcon). Nuclei were sorted on an MA900 cell sorter (Sony) equipped with a 100-μm microfluidic chip. Intact single nuclei were gated by FSC/BSC, and sfGFP-positive nuclei were collected using non-electroporated cortices as a negative control for background fluorescence.

### CUT&RUN

CUT&RUN was performed in two or three independent biological replicates per condition following the protocol of Skene et al.^55^ with minor modifications. Protein A-Protein G-MNase (pAG-MNase) was expressed in E. coli and purified in-house from the pAG/MNase plasmid (Addgene #123461) as previously described^56^. FANS-sorted nuclei (20,000 for histone modification profiling and 50,000 for other targets) were bound to concanavalin A-coated magnetic beads (BANG Lab, Cat# BP531-10), lightly fixed with 0.1% formaldehyde for 3 min at room temperature, and quenched with 125 mM glycine. Primary antibodies were added at the amounts shown in the key resources table. After MNase digestion and chromatin release, sonicated Drosophila melanogaster genomic DNA was added to the 2× Stop Buffer at a final concentration of 0.25 ng/mL as a spike-in control. After reverse cross-linking, DNA was purified by phenol-chloroform-isoamyl alcohol extraction and ethanol precipitation, followed by right-side size selection (0.5×) using Sera-Mag Select beads (Cytiva). For SUZ12 profiling, all wash and incubation buffers were adapted to physiological salt conditions as previously described^57^. Sequencing libraries were prepared using the NEBNext Ultra II DNA Library Prep Kit for Illumina (NEB, Cat# E7645) or KAPA EvoPrep Kit (Roche Cat# 10154039001) according to the manufacturer’s instructions, with a final double-sided size selection (0.5×–1.0×) using Sera-Mag Select beads to enrich for small fragments. CUT&RUN libraries were sequenced on an Illumina NextSeq2000 with paired-end reads.

### CUT&Tag

CUT&Tag was performed in three independent biological replicates using 50,000 FANS-sorted nuclei per sample, following a modified version of the previously described protocol^58^ ConA beads were prepared in-house as described previously^59^ by combining 10 µl of concanavalin A (2.3 mg/ml; Sigma, Cat# C2272) with 20 µl of Dynabeads MyOne Streptavidin T1 (Thermo Fisher Scientific, Cat# 65602). Cells were bound to these beads and incubated overnight at 4 °C in 300 µl of antibody buffer supplemented with primary antibody. Beads were washed in Dig-wash buffer, after which secondary antibody (guinea pig anti-rabbit IgG, 1:100; Rockland, ABIN101961) was applied for 1 h at room temperature. After a further Dig-wash step, samples were resuspended in Dig-300 buffer, and pAG-Tn5 (1:200; CST, Cat# 79561) was added for a 1 h incubation at room temperature. Beads were washed again in Dig-300 buffer, resuspended in tagmentation buffer, and held at 37 °C for 1 h to allow tagmentation. Tagmented DNA was recovered by phenol–chloroform–isoamyl alcohol extraction followed by ethanol precipitation. Libraries were generated with Q5 High-Fidelity 2X Master Mix (NEB, Cat# M0492S) following the manufacturer’s protocol, and small fragments were enriched by double-sided size selection (0.5×–1.0×) with Sera-Mag Select beads. Completed CUT&Tag libraries were sequenced on an Illumina NextSeq 2000 with paired-end reads. For FOS profiling, all wash and incubation buffers were adapted to physiological salt conditions as previously described^57^.

### Hi-C

In situ Hi-C was performed following the Hi-C 3.0 protocol⁵⁶ with minor modifications. E15 cortices were dissociated into single cells with Nerve Dispersion Solution (Wako, Cat# 291-78001) according to the manufacturer’s instructions, whereas P7 cortices were subjected to Dounce homogenization as described above. Dissociated cells and Dounce-homogenized nuclei were dual cross-linked with 1% formaldehyde for 10 min, followed by 3 mM disuccinimidyl glutarate (DSG; Thermo Fisher Scientific, Cat# 20593) for 40 min at room temperature, and quenched with 400 mM glycine. Cross-linked cells and nuclei were then immunostained with anti-PAX6 and anti-NeuN antibodies for 20 min at 4°C, and 100,000 PAX6⁺/NeuN⁻ cells from E15 and 100,000 NeuN⁺ nuclei from P7 were sorted per replicate at 4 °C and stored at −80 °C. Sorted cells and nuclei were permeabilized and digested overnight at 37 °C with DpnII (NEB, Cat# R0543). Digested ends were filled in with biotin-14-dATP (Thermo Fisher Scientific, Cat# 19524016), followed by in situ proximity ligation with T4 DNA ligase (NEB) at room temperature. After reverse cross-linking and DNA purification, ligated DNA was sonicated to fragments of approximately 300–500 bp using a Covaris focused-ultrasonicator, and biotinylated junctions were enriched with Dynabeads MyOne Streptavidin T1 (Thermo Fisher Scientific, Cat# 65602). Sequencing libraries were prepared on-bead using the KAPA EvoPlus v2 kit (Roche, Cat# 9420037001) according to the manufacturer’s instructions. Libraries were sequenced as paired-end reads on an Illumina NextSeq 2000.

### RNA-seq

Total RNA was extracted using RNAiso Plus (Takara Bio, Cat# 9109) according to the manufacturer’s instructions. For FANS-sorted nuclear RNA-seq (except for the Fig. 7 experiment) in SUN1-sfGFP expressing nuclei from neocortices, libraries were prepared using a modified FLASH-seq protocol^60^ with purified total RNA as input. dNTPs in the lysis/RT buffer were reduced to 3 mM, and betaine was omitted from this step; downstream reverse transcription (SuperScript IV), cDNA amplification (KAPA HiFi HotStart ReadyMix), TDE1 tagmentation, and index PCR followed the original protocol. For tissues harvested from E15 embryos and 8–9-week-old mice and for the experiment in Fig. 7, stranded RNA-seq libraries were prepared from total RNA using the SMARTer Stranded Total RNA-Seq Kit v3 – Pico Input Mammalian (Takara Bio, Cat# 634411) according to the manufacturer’s instructions. Libraries were sequenced on an Illumina NextSeq 2000 with paired-end reads.

### CUT&RUN and CUT&Tag data analysis

Paired-end reads were aligned to a hybrid reference genome comprising the mouse (mm10) and Drosophila melanogaster (dm6) genomes using Bowtie2^61^ (v2.4.5) with default settings, sorted and indexed with SAMtools^62^ (v1.14), deduplicated using Picard (v2.26.10), and filtered to remove reads overlapping the ENCODE mm10 blacklist^63^ using bedtools^64^ (v2.30.0). For spike-in normalization, a per-sample scaling factor was calculated as 10⁵ / N_dm6, where N_dm6 is the number of reads uniquely mapped to dm6, and spike-in-normalized BigWig tracks were generated using bamCoverage in deepTools^65^ (v3.5.1) with --extendReads --binSize 10, and the scaling factor passed via --scaleFactor. Peaks were called from spike-in-normalized BigWig files using LanceOtron^66^ (v1.0.8) with the options -t 4 -w 1000 for H3K27me3 at P7, -t 7 -w 1000 for H3K27me3 at E15, -t 15 -w 1500 for H3K4me3, and -t 4 -w 800 for SUZ12. Broad domains were called using SICER2^67^ (v1.0.3) against paired IgG controls with window size 500 bp, gap size 2500 bp, redundancy threshold 2, and FDR ≤ 0.05 for H3K27me3 at P7, -w 500 -g 3000 -fdr 0.01 for Lamin B1, -w 200 -g 400 -fdr 0.01 for H3K9me2, and -w 200 -g 5000 -fdr 0.01 for H3K9me3. H3K27me3 mega-domains were defined as SICER domains exceeding 0.5 Mb, which were obtained by merging domains called from the replicates and subsequently combining adjacent domains located within 100 kb of each other.

Peaks were annotated to nearby genes using HOMER^5^ (annotatePeaks.pl; v4.11), and those annotated as intergenic were excluded from subsequent analyses. Domains were annotated to gene bodies overlapped with the domains using bedtools intersect. Heatmaps and metaplots over features of interest were generated using computeMatrix and plotHeatmap / plotProfile from deepTools. Bins with signal in the top 0.005% were excluded from the BigWig files prior to generating metaplots to remove extreme outlier signals.

For genome coverage analysis, spike-in normalized H3K27me3 signal was calculated for each 5 kb bin. The bins from all time points within the same batch were pooled, and the top 20th percentile of the rolling mean values of H3K27me3 signal with a 50 kb sliding window was used as the threshold. Bins with a rolling mean value equal to or above this threshold were defined as H3K27me3^+^ regions at each time point. H3K27me3^+^ regions were annotated to genomic categories using the GenomicRanges^68^ and rtracklayer^69^ packages in R, with gene models from TxDb.Mmusculus.UCSC.mm10.knownGene (mm10). Promoters were defined as −2,000 to +200 bp around TSS, and gene bodies as the remaining gene regions after excluding promoters. The base pairs in each category were expressed as a percentage of total H3K27me3^+^ region length.

Clustering analyses were performed on spike-in–normalized bigWig files. Unless otherwise noted, k-means clustering was carried out using deepTools. For Fig. 5D, k-means clustering (k = 4) was performed in R using the H3K27me3, Lamin B1, H3K9me2, and mCH signals within H3K27me3 MDs. For Fig. 7H, FOS peaks called in KA-injected samples were used as reference regions, and k-means clustering of FOS signals from KA-injected samples overexpressing EV or JmjC was performed with fluff^70^ (v3.0.4; -C k -k 4 -e 5000 -g). Cluster 3 from this analysis was subsequently subclustered by k-means clustering of H3K27me3 signals from control (non–KA-injected) samples overexpressing EV or JmjC (-C k -k 2 -e 5000).

To assess transcriptional directionality, H3K4me3 peaks (grouped by k-means clustering in the H3K4me3/H3K27me3 heatmap) were annotated to genes with HOMER (annotatePeaks.pl, mm10) using the UCSC mm10 GTF. Intergenic peaks were excluded, and a unique gene list was generated per cluster. The sense-strand orientation of each gene was retrieved from the GTF, and the number of plus- and minus-strand genes was counted per cluster. Strand bias was tested for each cluster with a two-sided binomial test against the null expectation of equal strand proportions (P = 0.5), and p-values were corrected across clusters using the Benjamini–Hochberg method.

### RNA-seq data analyses

Raw reads were aligned to the mouse reference genome (mm10/GRCm38, UCSC) using STAR^71^ (v2.7.10a) with the following parameters: --outSAMtype BAM SortedByCoordinate --quantMode TranscriptomeSAM -- outFilterMultimapNmax 20 --alignSJoverhangMin 8 --alignSJDBoverhangMin 1 --outFilterMismatchNmax 999 -- alignIntronMin 20 --alignIntronMax 10000 --alignMatesGapMax 1000000 --outFilterScoreMinOverLread 0.3 -- outFilterMatchNminOverLread 0.3 --seedSearchStartLmax 12. Transcript-level quantification was performed using RSEM^72^ (v1.3.1) with options --paired-end --estimate-rspd --strandedness none --fragment-length-min 20 -- fragment-length-max 1000. Gene-level expected counts (rounded to integers) and TPM values were used for downstream analyses. Samples sequenced across multiple runs were combined by concatenating FASTQ files prior to alignment. Differential gene expression analysis was performed using sva^73^ (v3.52.0) and DESeq2^74^ (v1.44.0) in R (v4.4.2). Genes with an adjusted p value < 0.1 and |log₂ fold change| > 1 were considered differentially expressed.

### Hi-C data analyses

FASTQ files from three sequencing batches were concatenated for each sample and replicate. To ensure equal sequencing depth between E15 and P7 at each replicate, reads were downsampled to the smaller read count of the two samples using seqtk (v1.5, seed = 42). The downsampled FASTQ files from all four biological replicates were then merged for each sample. Hi-C data were processed using HiC-Pro^75^ (v3.0.0). Paired-end reads were aligned to the UCSC’s mm10 genome using Bowtie2 with the following options: --very-sensitive -L 30 --score-min L,-0.6,-0.2 --end-to-end for the global alignment step, and --very-sensitive -L 20 --score-min L,-0.6,-0.2 --end-to-end for the local alignment step, and non-duplicated valid interaction pairs were generated. Contact matrices were constructed at resolutions of 20, 40, 150, 500, and 1,000 kb, and normalized using iterative correction and eigenvector decomposition (ICE) with a maximum of 100 iterations, filtering out bins in the lowest 2% of coverage.

The HiC-Pro output contact matrices at 20 kb resolution were converted to cool format using cooler. Insulation scores were calculated from the 20 kb contact matrices using cooltools^76^ insulation (v0.7.1) with window sizes of 400 kb, excluding the two diagonals nearest to the main diagonal (--ignore-diags 2). The log2-transformed insulation scores were exported as BigWig files for visualization, excluding bad bins and bins with missing values. Genome tracks were generated with pyGenomeTracks^77,78^ (v3.9).

TAD boundaries were defined by applying an Otsu threshold to boundary strength scores at a 400 kb window, using the average threshold calculated from E15 and P7. TADs were classified as H3K27me3-MD TADs if at least 80% of their length overlapped with H3K27me3+ regions, as determined using bedtools coverage. To compare Hi-C contact frequencies between H3K27me3-MD and -non-MD TADs, observed/expected (OBS/EXP) contact scores were calculated for each bin pair at P7 using the ICE-normalized contact matrix. Intra- or inter-TAD contact frequency was summarized as the mean OBS/EXP value across all bin pairs within or in-between the TADs. The two nearest diagonals and bins with missing values were excluded after ICE normalization.

Aggregate TAD Analysis (ATA) was performed for H3K27me3-MD and -non-MD TADs at E15 and P7. TADs where more than 50% of values were missing after normalization were excluded. Each matrix was resized to 99 × 99 bins using bilinear interpolation, and pileup matrices were computed by taking the nanmean across all TADs. Differential ATA was calculated as the difference in log2(OBS/EXP) between H3K27me3-MD and -non-MD TADs for each time point.

### WGBS data analyses

Trimmed reads were aligned to the UCSC’s mm10 genome using Bismark^79^ (v0.24.2) with Bowtie2 (-N 1 -L 20). Duplicate reads were removed using deduplicate_bismark, and methylation information was extracted using bismark_methylation_extractor with the first 2 bases of each read ignored. CHG and CHH context files were merged to generate a single non-CpG methylation (mCH) file, and methylation values were averaged over 100 bp bins using bedtools map.

### Motif enrichment analysis

Motif enrichment analysis was performed on 363 regions using findMotifsGenome.pl from the HOMER suite (v4.11), with the -size 100 option and a GC content-matched genomic background. Only de novo motifs are reported.

### Gene Ontology analysis

Gene Ontology (GO) enrichment analysis was performed using Enrichr^80^ (https://maayanlab.cloud/Enrichr/). The “GO Biological Process 2025/2026” libraries under the Ontologies category were used to identify enriched terms for each gene list. Significant terms were defined as those with an adjusted P value < 0.05.

### Whole body gene expression analysis

Baseline gene expression data (TPM) were downloaded from the EMBL-EBI Expression Atlas (accession: E-MTAB-3579), originally derived from the RIKEN FANTOM5 project (Mus musculus)^81–83^, and expression values from juvenile tissues were extracted. To quantify tissue expression bias toward non-brain tissues, a Dz score was calculated for each gene. Log2-transformed TPM+1 values were converted to z-scores across all tissues for each gene. Tissues were grouped into brain (cerebellum, cerebral cortex, diencephalon, and medulla oblongata) and non-brain (adrenal gland, bone, epididymis/seminiferous tubule, heart, intestine, kidney, liver, lung, muscle, pancreas, spleen, stomach, testis, and thymus) categories. The Dz score was defined as the difference between the maximum z-score among non-brain tissues and the maximum z-score among brain tissues (Dz = NBmaxZ − BmaxZ), where positive values indicate a non-brain expression peak and negative values indicate a brain expression peak. Dz scores of H3K27me3 mega-domain genes were compared with neuronal genes (annotated with the GO term “chemical synaptic transmission” (GO:0007268, Mus musculus, annotation class label: chemical synaptic transmission), 116 genes), lysosomal genes (annotated with the GO term “regulation of lysosome organization” (GO:1905671, Mus musculus), 14 genes) and all genes.

## QUANTIFICATION AND STATISTICAL ANALYSIS

Statistical analyses for all experiments were performed in R or Python. Numbers of experimental replicates, *P* values and the tests can be found in the figure legends and Methods. The box denotes the interquartile range (IQR), and the whiskers denote the rest of the data distribution.

---

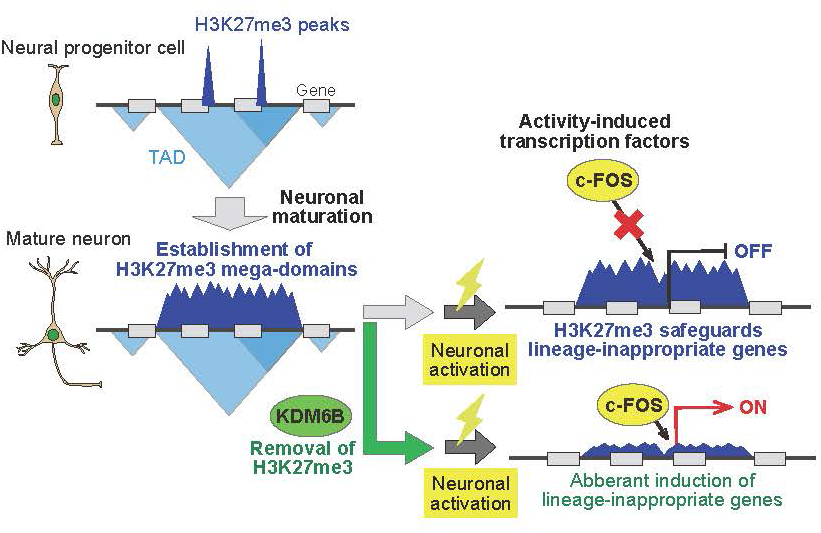

**Supplemental Figure 1.**
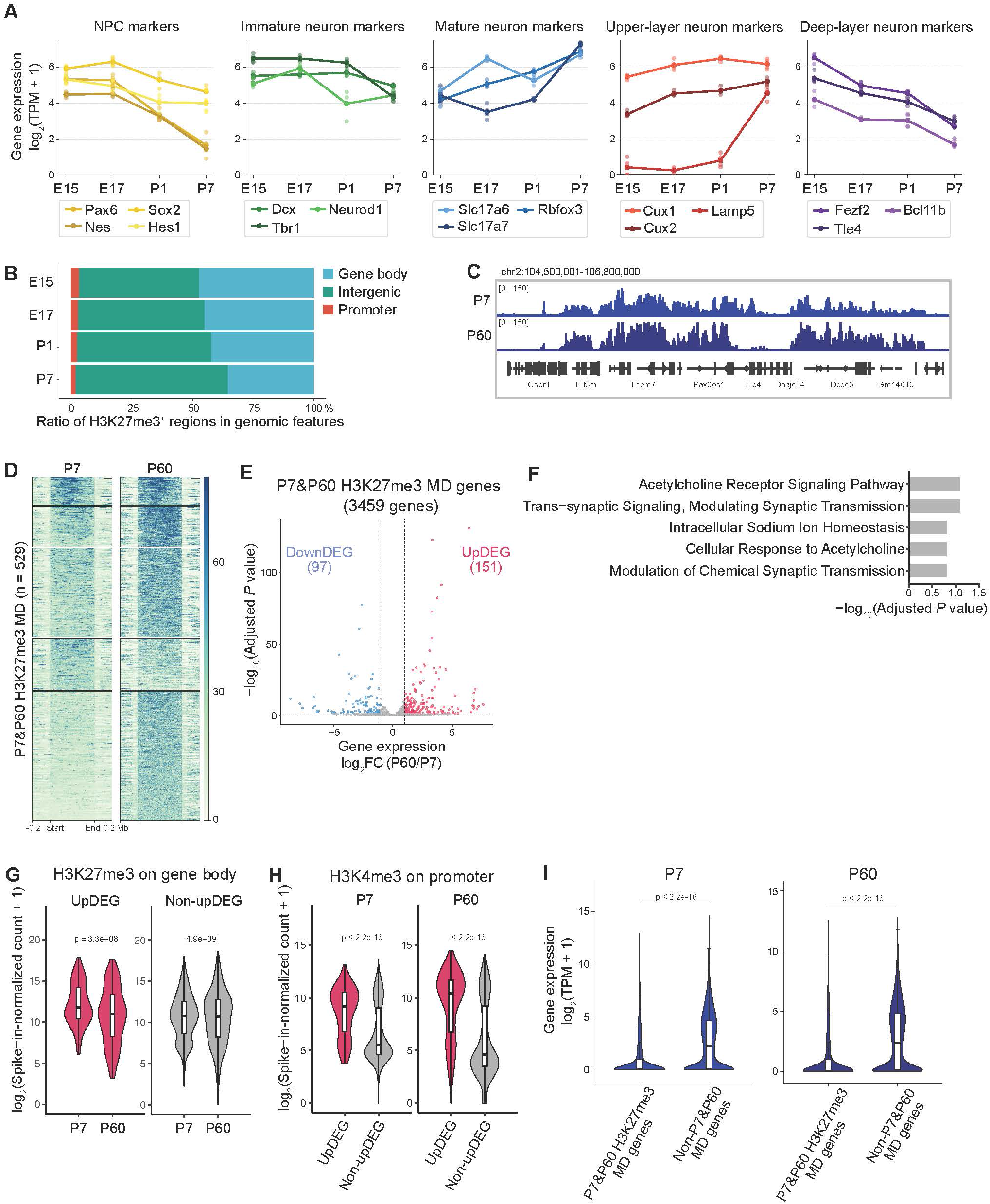
Characterization of the SUN1-sfGFP-labeled neuronal lineage and continued reorganization of H3K27me3 domains between P7 and P60 (A) Line plots showing the expression levels of neuronal cell-type marker genes in SUN1-sfGFP^+^ nuclei across developmental stages (n = 4 biological replicates per stage). Marker genes include neural progenitor cells (*Pax6, Sox2, Nes* and *Hes1*), immature neurons (*Dcx, Neurod1* and *Tbr1*), pan-mature neurons (*Slc17a6, Slc17a7* and *Rbfox3*), upper-layer excitatory neurons (*Cux1, Cux2* and *Lamp5*) and deep-layer excitatory neurons (*Fezf2, Tle4* and *Bcl11b*). The results confirmed the expected progression of neuronal differentiation and maturation. (n = 4 biological replicates per stage, 50,000 nuclei per sample). (B) Bar graph showing the relative genomic distribution of H3K27me3^+^ regions at each developmental stage. Regions were annotated as promoters, gene bodies, or intergenic regions, with promoters defined as regions from 2 kb upstream to 200 bp downstream of the TSS. (C) Genome browser snapshot showing spike-in-normalized H3K27me3 signals at representative H3K27me3 target gene loci in P7 and P60. (D) Heatmap showing spike-in-normalized H3K27me3 signals at merged H3K27me3 MDs at P7 and P60. Merged MDs were defined as the union of H3K27me3 MDs identified at P7 and P60 (n = 529 regions; 4 biological replicates per stage) and clustered by k-means clustering (k = 5). (E) Volcano plot showing differential expression of H3K27me3 MD genes between P7 and P60. Genes with adjusted P < 0.05 and |log_2_ fold change| ≥ 1 were classified as upregulated DEGs (UpDEGs) or downregulated DEGs (DownDEGs) and are highlighted. (F) Bar graph showing selected GO Biological Process 2026 terms enriched among UpDEGs identified in Figure S1E. GO enrichment analysis was performed using 151 upregulated genes associated with merged P7&P60 H3K27me3 MDs. Bars indicate -log10-transformed adjusted p-values. (G) Violin plots showing gene-body H3K27me3 signal intensity at P7 and P60 for UpDEGs and non-UpDEGs identified in Figure S1E. (H) Violin plots showing the spike-in-normalized H3K4me3 signal in the promoters of UpDEG and non-UpDEG H3K27me3 MD genes at P7 and P60. Promoters were defined as regions from 1 kb upstream to 100 bp downstream of the TSS. (I) Violin plot showing the expression levels of merged H3K27me3 MD genes at each developmental stage (P7 and P60). Merged MD genes were defined as the union of H3K27me3 MD genes identified at P7 and P60 (n = 3,459 genes) and other genes were defined as non-MD genes (n = 20951). (4 biological replicates per stage, 50,000 nuclei per sample). Statistical significance was assessed by a Mann-Whitney U test, and p-values are indicated.

**Supplemental Figure 2.**
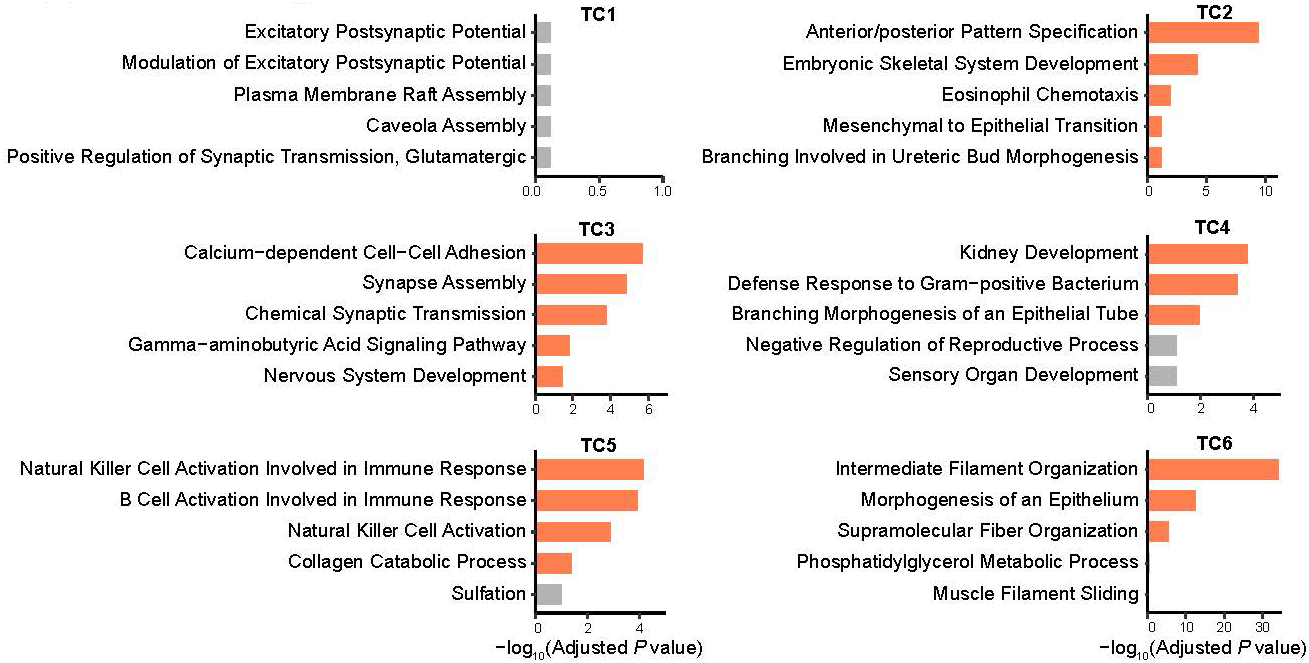
Gene Ontology analysis of H3K27me3 mega-domain clusters in the five tissues Bar graphs showing selected GO Biological Process 2026 terms enriched among genes associated with each cluster defined in Figure 2E. Genes were assigned to a cluster if their gene bodies were located within a merged H3K27me3 MD region belonging to that cluster. Gene counts per cluster were as follows: TC1, 1,709; TC2, 642; TC3, 967; TC4, 1,377; TC5, 971; TC6, 1,575. Orange bars indicate significantly enriched terms, defined by Benjamini–Hochberg-adjusted p < 0.05; gray bars indicate non-significant terms shown for context. Bars represent −log10-transformed adjusted p-values. X-axis scales differ between clusters.

**Supplemental Figure 3.**
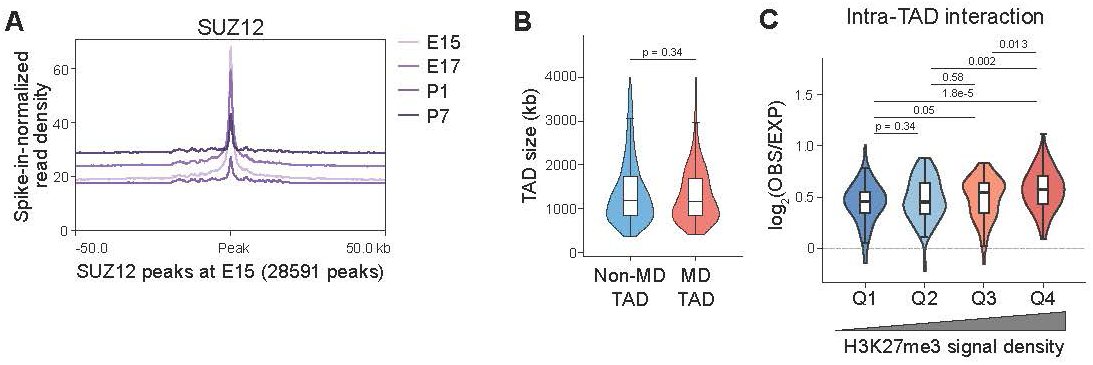
SUZ12 occupancy remains restricted during maturation, and H3K27me3 levels correlate with intra-TAD interaction strength (A) Metaplot showing average spike-in-normalized SUZ12 signal profiles at E15.5, E17, P1, and P7 centered on SUZ12 peaks at E15.5 (n = 28,591 peaks; ±50 kb flanks). (B) Violin plot showing TAD size distributions of MD TADs and non-MD TADs at P7. MD and non-MD TADs were classified as in Figure 3F. P-value was calculated using a Mann-Whitney U test. Note that TAD size did not differ between the two groups, indicating that the increased interaction frequency cannot be explained by domain length alone. (C) Violin plot showing intra-TAD contact frequencies of MD TADs at P7 stratified into quartiles (Q1–Q4) based on H3K27me3 signal density within each TAD, quantified as the mean H3K27me3 signal per bin. P-values were calculated using a Mann-Whitney U test followed by Benjamini-Hochberg correction.

**Supplemental Figure 4.**
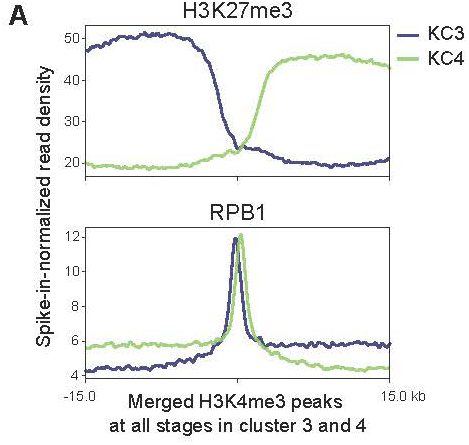
RNA polymerase II and H3K27me3 distribution at H3K4me3 peaks flanking H3K27me3 mega-domains (A) Metaplot showing the average spike-in-normalized H3K27me3 and RPB1 signal profiles at P7 around merged H3K4me3 peaks in cluster 3 and cluster 4 defined in Fig. 4E. Merged H3K4me3 peaks were defined as the union of H3K4me3 peaks identified across all four developmental stages. Cluster 3 contains 3,353 peaks, and cluster 4 contains 3,819 peaks. Signals are shown across ±15-kb flanking regions.

**Supplemental Figure 5.**
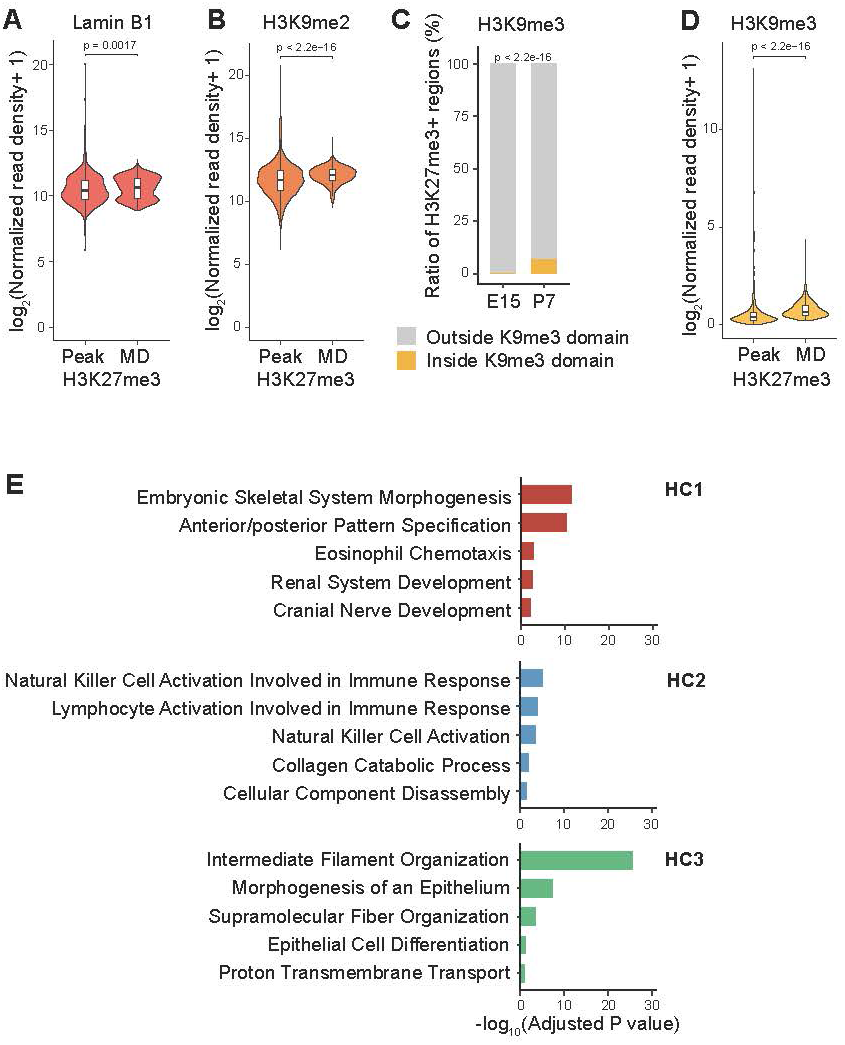
H3K27me3 mega-domains are enriched for heterochromatin-associated marks (A, B, D) Violin plots showing spike-in-normalized Lamin B1 (A), H3K9me2 (B) and H3K9me3 (D) read densities at P7 H3K27me3 peaks and P7-defined H3K27me3 MDs. P-values were calculated using a Mann–Whitney U test. Lamin B1, H3K9me2, and H3K9me3 profiles were generated from n = 2 biological replicates at P7. (C) Bar graph showing the proportion of H3K27me3^+^ regions located within H3K9me3-enriched domains (K9me3 domains) at E15 and P7. H3K9me3 domains were defined from H3K9me3 profiles generated from n = 2 biological replicates per stage. Statistical significance was assessed by Fisher’s exact test, and p-values are indicated. (E) Bar graph showing selected enriched categories (GO Biological Process 2026) for the genes included in each H3K27me3 mega-domain cluster as calculated by Enrichr. The genes in cluster 4 were not significantly enriched in any GO term.

**Supplemental Figure 6.**
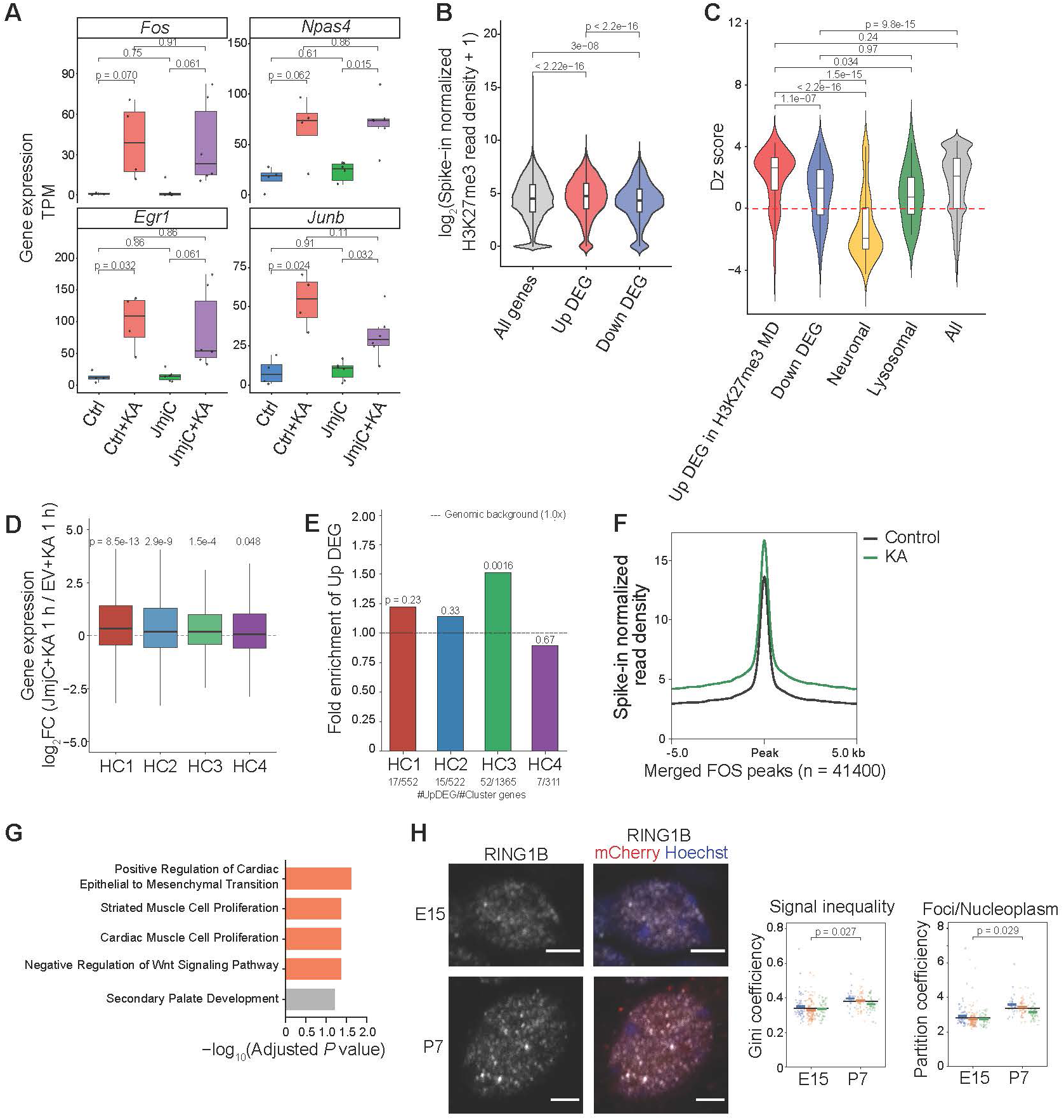
Activity-induced derepression of lineage-inappropriate genes upon H3K27me3 depletion (A) Box plots showing gene expression levels of the immediate early genes Fos, Npas4, Egr1, and Junb at P8 under four experimental conditions: control, KA-injected control, JmjC-expressing, and KA-injected JmjC-expressing samples. (B) Violin plot showing H3K27me3 read density over gene bodies for all genes, upregulated DEGs, and downregulated DEGs identified in KA-injected JmjC-expressing samples relative to KA-injected empty vector (EV) control samples. Read density was normalized by gene length and spike-in ratio. Statistical significance was assessed using a Mann–Whitney U test followed by Benjamini–Hochberg correction; adjusted p-values are indicated. (C) Violin plot showing the distribution of Dz scores for the five indicated gene categories, including DEGs identified in KA-injected JmjC-expressing samples relative to KA-injected empty vector (EV) control samples. Dz scores were calculated as described in Figure 1K: Dz score = maximum Z-score across non-brain tissues − maximum Z-score across brain tissues. Positive and negative Dz scores indicate preferential expression in non-brain and brain tissues, respectively. (D) Box plot showing log2 fold changes in gene expression for genes in the four clusters defined in Figure 5D in KA-injected JmjC-expressing samples relative to KA-injected empty vector (EV) control samples 1 h after KA injection. Gene numbers per cluster were as follows: HC1, 1,263 genes; HC2, 1,186 genes; HC3, 2,484 genes; HC4, 1,133 genes. Statistical significance for positive shifts from zero was assessed using a one-sided one-sample Wilcoxon signed-rank test; p-values are indicated above each box. (E) Bar graph showing fold enrichment of upregulated DEGs identified in KA-injected JmjC-expressing samples relative to KA-injected empty vector (EV) control samples within each cluster defined in Figure 5D. For each cluster, fold enrichment was calculated as the ratio of the proportion of upregulated DEGs within that cluster to the proportion of upregulated DEGs among all expressed genes. Statistical significance was assessed using a one-sided Fisher’s exact test to determine whether upregulated DEGs were enriched within each cluster relative to the background rate. (F) Metaplot showing average spike-in-normalized FOS signal profiles in control samples and KA injected samples at P8 centered on merged FOS peaks in empty vector (EV) control samples and KA injected samples (n = 41,400 peaks; ±5 kb flanks). (G) Bar graph showing p-values of enrichment of selected GO Biological Process 2026 terms. GO enrichment analysis was performed using 413 H3K27me3-MD genes on FC4 cluster. Bars indicate -log10-transformed adjusted p-values. (H) Representative images showing the signals of RING1B staining, mCherry, and Hoechst in an mCherry-labeled cell of the neocortex from E15 and P7 mice. The right plots show the statistical comparisons. The left plot shows the Gini coefficient of the voxel intensities indicating the inequality of the signal intensity inside a nucleus. The right plot shows the partition coefficient, defined as the ratio of the mean intensity of condensate voxels to the mean intensity of the remaining nucleoplasmic voxels. Individual nuclei and the mean value of each biological replicate (animal) were displayed in different colors, and per-animal means were compared between E15 and P7 by two-sided Welch’s t test (n = 3 animals per group; E15, 223 nuclei; P7, 145 nuclei in total). Scale bar, 3 µm.

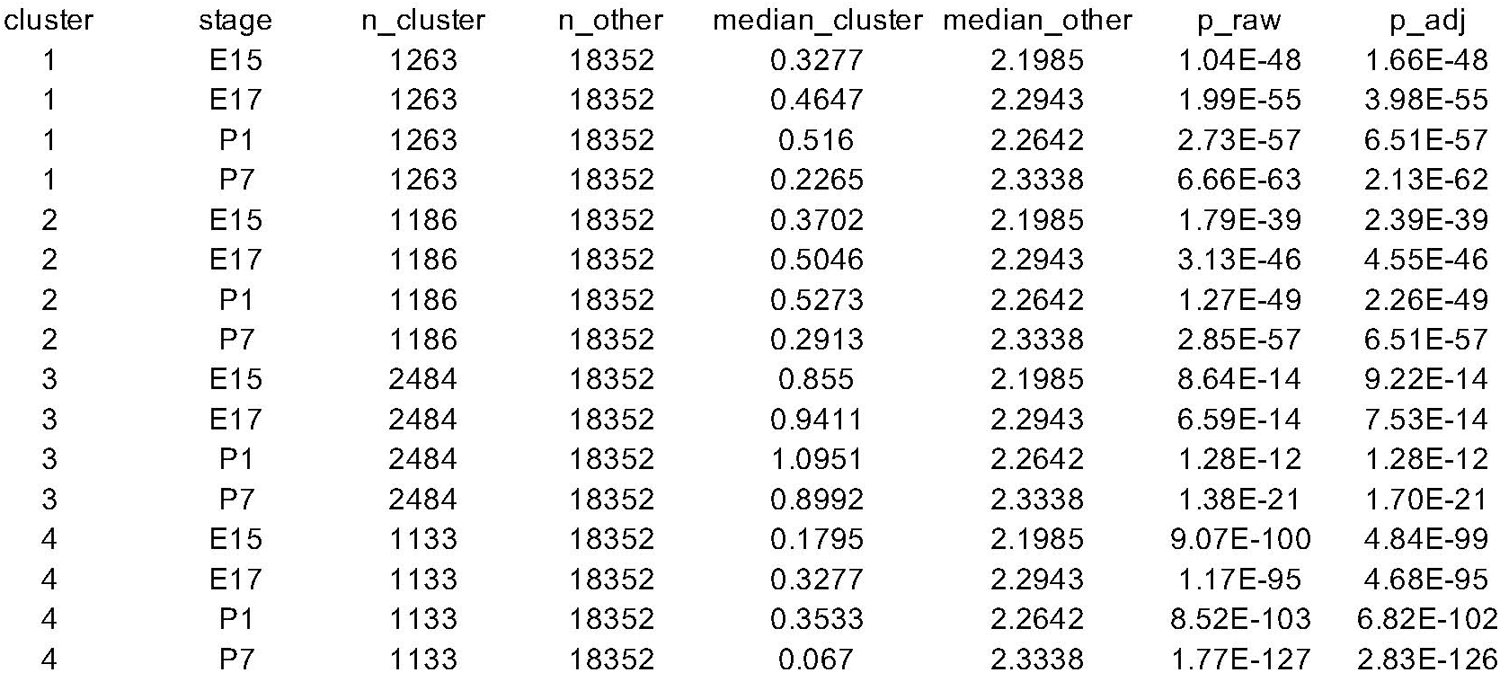

